# Evolutionary impact of size-selective harvesting on shoaling behavior: Individual-level mechanisms and possible consequences for natural and fishing mortality

**DOI:** 10.1101/809442

**Authors:** Valerio Sbragaglia, Pascal P. Klamser, Pawel Romanczuk, Robert Arlinghaus

**Author notes:** V.S. and P.P.K. contributed equally to this work. **Correspondence:** experimental data, models and simulations.

## Abstract

Intensive and size-selective harvesting is an evolutionary driver of life-history as well as individual behavioral traits. Yet, whether and to what degree harvesting modifies the collective behavior of exploited species is largely unknown. We present a multi-generation harvest selection experiment with zebrafish (*Danio rerio*) as a model species to understand the effects of size-selective harvesting on shoaling behavior. The experimental system is based on a large-harvested (typical of most wild capture fisheries targeting larger size classes) and small-harvested (typical of specialized fisheries and gape-limited predators targeting smaller size classes) selection lines. By combining high resolution tracking of fish behavior with computational agent-based modeling we show that shoal cohesion changed in the direction expected by a trade-off between vigilance and the use of social cues. In particular, we document a decrease of vigilance in the small-harvested line, which was linked to an increase in the attention to social cues, favoring more cohesive shoals. Opposing outcomes were found for the large-harvested line, which formed less cohesive shoals. Using the agent-based model we outline possible consequences of changes is shoaling behavior for both fishing and natural mortality. The changes in shoaling induced by large size-selective harvesting may decrease fishing mortality, but increase mortality by natural predators. Our work suggests an insofar overlooked evolutionary mechanism by which size-selective harvesting can affect mortality and in turn population dynamics of exploited fish.

## Introduction

Size-selective mortality is an important evolutionary driver in fishes that typically suffer greater natural mortality in juvenile than adult stages (1, 2). Human activities, specifically capture fisheries, profoundly alter the natural mortality-based fitness landscape by increasing adult mortality to levels hardly ever experienced in the evolutionary history of most fish populations (3). Size-selective mortality imposed by fishing can foster adaptive responses in terms of life-history as well as physiological and behavioral traits (4–7). Fishing can also affect shoaling tendency and collective behavior of fish (8, 9), with potential consequences for population dynamics and food-web functioning (10). However, the mechanisms through which size-selective harvesting affects shoaling behavior are largely unexplored.

Shoaling behavior has strong adaptive value in many fish species, reducing predation risk, increasing foraging efficiency, and reducing energetic costs (11–13). For example, a classic expectation is that fish increase shoal cohesion in the presence of predation risk because more cohesive, denser, and coordinated groups can more effectively confound predators and consequently reduce their predation efficiency (14). Shoaling behavior is complex and can be simultaneously influenced by several variables including predation risk (15), food availability (16), and light levels (17). In particular, the trade-off between vigilance (i.e. the frequency of environmental scans) and time spent feeding is a crucial aspect in determining shoaling behavior in fishes (15, 18). However, because coordinated movement demands attention to social cues, there is also a trade-off between shoaling and vigilance. For example, in three-spined sticklebacks, *Gasterosteus aculeatus*, more aligned individuals respond slower to an external cue (19), and in herring, *Clupea harengus*, solitary individuals respond faster to an external cue than shoaling individuals (20). Therefore, an increase of vigilance could weaken social coordination and hence shoal cohesion. Vigilance can also be affected by group size (21–23) and social behavior (e.g. territorial aggression; 24). Shoaling behavior is a key driver of catchability in fisheries (25), thus changes in shoaling will affect fisheries outcomes. Considering that collective behavior has shown rapid evolution in response to artificial selection in guppies, *Poecilia reticulata* (26), size-selective harvesting could move the fitness landscape of shoaling behavior away from natural optima (3). While a corresponding adaptation to the new fitness landscape is likely to reduce exposure of fish to harvesting, it might constitute a novel pressure to exploited populations by increasing natural mortality.

The collective mechanisms governing shoaling behavior in fish and other animals can be explained by simple interaction rules among individuals that are influenced by their respective neighbours (27–29). Major group-level behavioral transitions can be triggered by minor changes in social interactions at the individual level (30). Therefore, size-selective harvesting could affect shoaling behavior by changing individual behavioral traits that influence the interaction rules that drive group behavior. However, how shoal cohesion is influenced by the trade-off between vigilance and the use of social cues, or by directly responding to perceived risk, is to our knowledge largely unexplored in general and in response to size-selective harvesting more specifically. In this context, due to non-trivial connections between individual-level and group-level behavior, it is important to complement experimental data with agent-based models, to 1) identify emergent effects due to self-organized collective behavior not directly “encoded” in individual behavior (30–32) and 2) to investigate the consequences and mechanisms of individual-level adaptations on the dynamical behavior of the group (33).

We take advantage of a long-term artificial selection experiment of zebrafish *Danio rerio* as a model shoaling species to understand the effects of size-selective harvesting on shoaling behavior and test potential consequences for natural and fishing mortality. Zebrafish is a small-bodied non-obligated schooling species (34). Research on the behaviour of zebrafish in the wild suggests that zebrafish also regularly shoal in rivers and ponds (35, 36), but environmental conditions can affect such behavior. For example, zebrafish from slow-flowing rivers form smaller and less cohesive groups than zebrafish from fast-flowing rivers, and predation risk is suggested to increase shoal cohesion both in the wild (36) and laboratory (37). Swimming in groups can also increase food exploitation in zebrafish, collectively emphasizing the adaptive value that shoaling has for this species (38). In terms of swimming behavior, zebrafish exhibits a burst-coast swimming pattern that is characterized by an “active” and “passive” mode (39); specifically, during the active mode zebrafish are sensitive to the swimming patterns of conspecifics (39), and consequently to other stimuli related to perceived risk and vigilance. Overall, zebrafish is a suitable candidate model to test how size-selective harvesting evolutionary affect the trade-off between vigilance and social information and its relative impact on shoal cohesion.

In our experimental system, artificial selection has been imposed on wild-collected zebrafish populations (5), therefore the mechanisms investigated here could be relevant for wild shoaling fish populations exposed to intense directional selection. In particular, the experimental lines were subjected to five generations of opposing size-selective harvesting (5): large-harvested line (larger individuals harvested; a common scenario in many fisheries worldwide and in presence of predators where large individuals are selectively preyed upon), small-harvested line (smaller individuals harvested; a possible scenarios in specific fisheries or in the presence of gape-limited predators that preferentially feed on the smaller size classes) and random harvested line (control). Previous results revealed substantial changes in life history, body size, gene expression, allele frequencies as well as individual and group behavioral traits (5, 7, 40–44). In particular, the small-harvested zebrafish selection line increased boldness during feeding and after simulated predator attacks, while the large-harvested showed opposed but weaker effects on boldness compared to controls (7). These experimental findings largely agree with a recent theoretical model that predicted that large size-selective harvesting could either foster bold or shy fish, depending on the strength of size-selection and correlations of behavior and growth rate (6).

We hypothesized that the evolutionary effect of size-selective harvesting on shoaling behavior is based on the trade-off between vigilance and the use of social cues (12), where an increase of individual vigilance is linked to a decrease of attention to social cues with consequences in emerging shoal cohesion. We assumed that the increase of boldness in the small-harvested line is linked to a decrease of vigilance and consequently fosters an increase in shoal cohesion (vice versa for the large-harvested line). Furthermore, we used a computational modelling approach to disentangle the mechanistic process driving the possible differences in shoal cohesion among the selection lines (Fig. 1). Specifically, we present a novel burst-coast, agent-based model to link individual-level traits (e.g., vigilance modeled as burst rate using environmental vs. social cues) to group-level emergent properties such as shoal cohesion (Fig. 1). We used high resolution tracking of individual fish within groups to calibrate and fit the model to experimental data. This allowed to asses at what extent the use of environmental vs. social cues drives changes in shoal cohesion (Fig. 1). Lastly, with the model representations of the selection lines, we simulated shoaling behavior in the presence of a natural predator or fishing gear to explore how size-selective harvesting could affect mortality in natural and fishing contexts (Fig. 1). In summary, our paper presents novel experimental data on the possible evolutionary effect of size-selective harvesting on shoaling behavior, theory and modelling to mechanistically explain the underlying processes driving the observed changes, and simulated consequences to predict possible effects on fishing and natural mortality.

**Fig.1.**
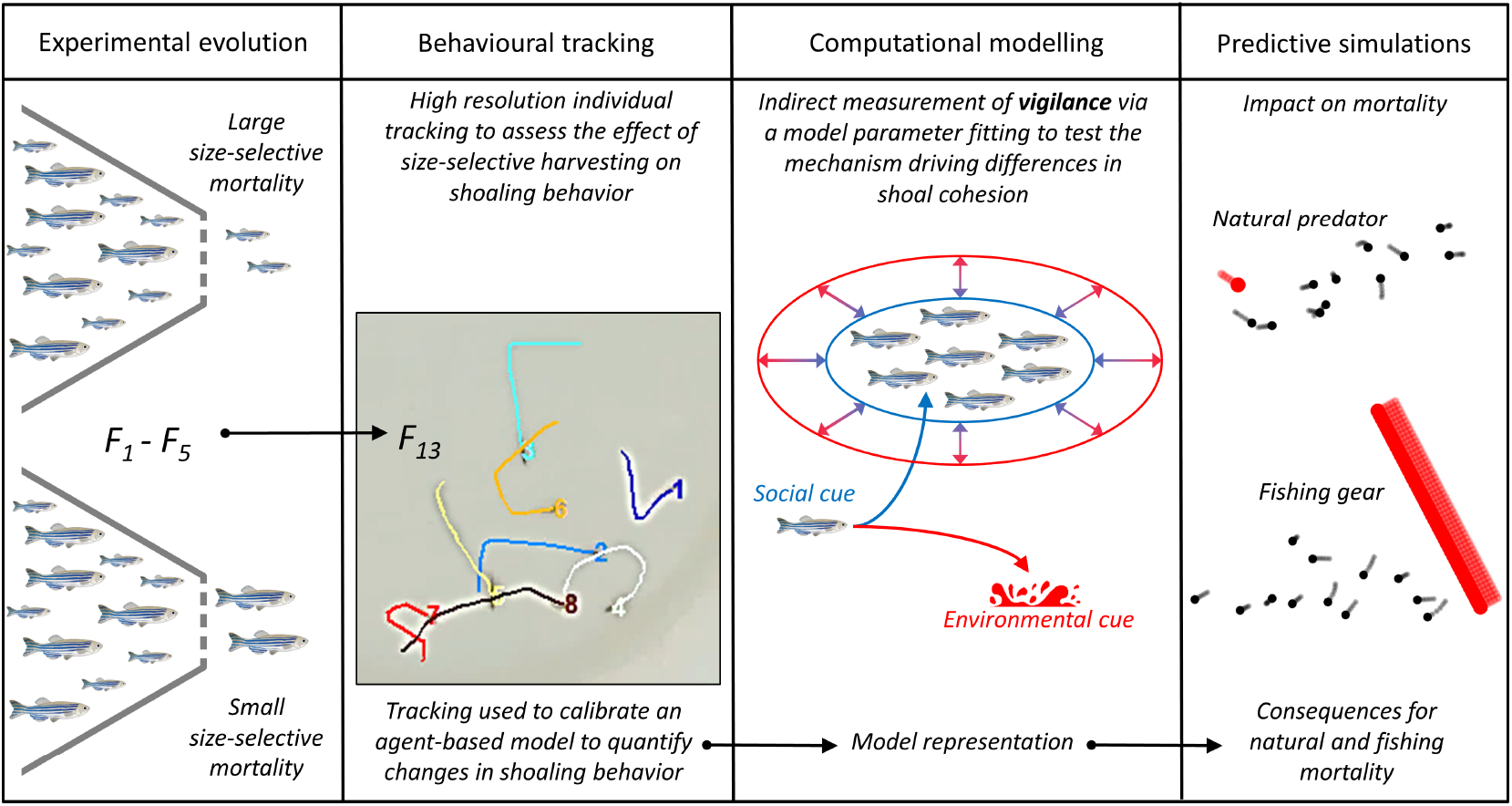
The integrative research framework used here is presented according to four major blocks. Experimental evolution in wild-collected zebrafish was used to test the effect of size-selective harvesting on shoaling behavior (selection occurred during the first five generations and then stopped for eight additional generations; see methods). Behavioral tracking of individual fish within shoals was performed to measure individual (e.g., speed) as well as group (e.g., shoal cohesion) variables among size-selective harvesting treatments. Computational modelling was used to underpin the mechanisms governing changes in shoaling behavior. An agent-based model was calibrated using the experimental results of behavioral tracking; next, we obtained an indirect measurement of vigilance using a model parameter fitting. Finally, the model was used to simulate different scenarios with the goal to assess how changes in shoaling behavior in response to size-selective harvesting could affect natural and fishing mortality.

## Methods

### A. Selection lines

Our study system consisted of wild-collected zebrafish from West Bengal in India (for more details see 41, 45). The parental wild-collected population was experimentally harvested over five generations (5) by exposing them to strong directional selection (a harvest rate of 75 percent per-generation) for either small body size (large-harvested line) or large body size (small-harvested line). A third line was harvested randomly with respect to size and used as a control (random-harvested or control line). Each selection line was replicated twice, which adds up to six selection lines. After five generations the harvesting was stopped for eight further generations until F13 – a time when the present experimental trials took place. At F9 the large-harvested line evolved a smaller adult length and weight and higher relative fecundity compared to controls (5). By contrast, the small-harvested line showed reduced reproductive investment and no change in adult length compared to the control line (5). Both size-selected lines matured smaller and at younger age than the control line (for more details see 5). These differences persisted until F13 (42, 43). Overall, the size-selective harvesting fostered changes in energy allocation patterns, with the large-harvested line investing early and intensively into reproduction, and the small-harvested line investing early but at low intensity into gonad production and more intensively in somatic growth.

### B. Experimental design

We randomly selected 6 groups of 8 juvenile zebrafish at 30 days post fertilization for each of the selection lines, which consisted of at about 450 zebrafish per selection line (N=12 groups per selection treatment). The 36 groups were housed in three-liter rearing boxes in random order on shelves of the same holding system. Throughout the experiment zebrafish were fed ad libitum with dry food (TetraMin, Tetra), the water temperature was maintained at 26±0.5 °C and photoperiod at 12:12 h light-darkness cycle (light on/off at 07:00 and 19:00, respectively). All trials were run between 09:00 and 14:00. We started the experimental trials two hours after light-on to avoid measuring mating behavior that usually occurs in the first hours after light-on (46). The movement of the fish from the rearing boxes to the experimental tanks was conducted by gently pouring them together with the water from the stocking box. The experimental protocols were approved by the Spanish National Committee and the Committee of the University of Murcia on Ethics and Animal Welfare (CEEA-OH 491/2019).

### C. Shoaling trial

We performed the shoaling trials with the groups of 8 zebrafish at two time points (150 and 190 days post fertilization when all individuals were adults). Shoaling was measured in a white round arena (diameter of 49 cm) with 10 cm of water. The arena was placed on a table behind a white curtain to minimize disturbance to the fish during the experimental trials. We recorded behavior using a webcam (resolution: 1920 x 1080 pixels; frame rate: 30 frames per second) from about 1m above the arena. Zebrafish were introduced in the experimental arena and left undisturbed for 25 min before starting the experimental trial. Video recording lasted 5 min and after that we measured the standard length of each fish on a petri dish with millimeter paper anesthetizing the fish using a clove oil dilution in ethanol and water. First, in order to test repeatability of shoaling behavior, we analyzed the video recordings at both time points (150 and 190 days post fertilization) with automated behavioral tracking (EthoVision XT 9, Noldus Information Technologies Inc.; www.noldus.com). Then, in order to get individual trajectories, we also analyzed the video recordings at 190 days post fertilization with the idTracker software. The software extracts specific characteristics of each individual and uses them to identify each fish without tagging (47).

### D. Data analysis

We first extract measures of mean inter-individual distance (i.e., the average distance of a given fish from all the other fish in the arena) from the output of the group tracking (150 and 190 days post fertilization), where we were not able to distinguish the individual fish within the shoal. Then, we characterized the output of individual tracking of shoaling behavior at 190 days post fertilization according to three group-level variables: inter-individual distance, nearest-neighbor distance (i.e., the distance from a focal fish to its closest neighbor averaged over all fish, NND), and polarization (i.e, the absolute value of the mean heading direction). We only considered the frames where all the shoal members were correctly detected (30 out of the 36 trials were used; N = 10 for each line treatment). We also calculated individual-level variables such as speed and burst rate. In this case, the output of the behavioral tracking was filtered by excluding frames with a probability of correctly identifying an individual below 0.85. This increased the tracking quality, but reduced the number of trials used in the individual analysis (18 out of the 36 trials were used; N = 6 for each line treatment). The modelling approach explained below focused on burst and coast phases, therefore individual identification was vital because properties of the burst phase, as its duration and rate, can be corrupted by tracking inaccuracy. Assuming that the remaining inaccuracy was random, we used a moving average with a Gaussian kernel to correct it (Figs. S7, S6).

### E. Statistical approach for behavioral data

The measures of mean inter-individual distance derived from the group behavioral tracking at 150 and 190 days post fertilization were transformed by finding the exponent (*λ*) which makes the values of the response variable as normally distributed as possible with a simple power transformation. Next, we calculated the adjusted repeatability (i.e., after controlling for fixed effects of selection lines) according to Nakagawa and Schielzeth (48). The replicates of the selection lines were used as additional random intercepts in the model. The results of the individual tracking at 190 days post fertilization were analyzed as follows. Group (inter-individual distance; nearest-neighbor distance, and polarization) and individual (body size, speed and burst rate) measurements were (*λ*) transformed as reported above. Then, response variables were modelled using linear mixed effects models with selection lines as fixed effect and selection line replicate as random intercepts. Likelihood ratio tests were used to assess the significant effect of selection lines as fixed effect (95 % confidence interval). Considering that zebrafish individual behavior is correlated with size (49) and that the size-selective treatments modified the size at age of the selection lines (5), it is conceivable that individual size differences among the selection lines could have masked harvesting-induced changes in terms of individual behavior. We thus estimated the effect of size (standard length) as a covariate against individual behavior. We implemented a second set of models with a size-matched dataset. The response variables were modelled by using linear mixed models and implementing all possible models (i.e., four) with individual body length as covariate, selection lines as fixed effect and selection line replicate as random intercepts. Likelihood ratio tests were used to assess the significant effect of selection lines and body length (95 % confidence interval). All statistical analysis were implemented using R 3.5.0 (https://www.R-project.org/).

### F. Shoaling model (burst and coast model)

To unravel the mechanistic underpinning of our experimental results, we implemented a burst-coast-agent-based model (Fig. 2; Movie S1; Tab. 1). We built this model with the purpose to represent the experimental data and to assess at what extent the use of environmental vs. social cues drives changes in shoal cohesion (Fig. 1). Therefore, the model is not intended to represent evolutionary mechanisms, but rather to investigate the mechanisms driving the experimental results linking individual-level traits (e.g., swimming speed and burst rate using environmental vs. social cues) to group-level emergent properties such as shoal cohesion (Fig. 1). In order to do that, we reproduced the burst-coast swimming typical of zebrafish by a burst phase, initiated at a rate *γ*, in which a fish *i* accelerates along a constant force 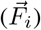 for the duration of the burst (*t_b_*). If the fish does not burst, it coasts (i.e. its acceleration force is zero 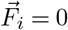) by decelerating due to friction (friction coefficient *β*) and keeps its heading direction (*ê*_||,*i*_). This behavior corresponds to the changes in position 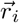 and velocity 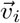 according to

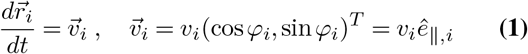

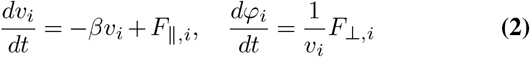

with *v_i_* and *φ_i_* being the fish speed and its heading direction, which are altered by the force components parallel *F*_||,*i*_ and perpendicular *F*_⊥,*i*_ to the heading direction *ê*_||,*i*_, respectively (50).

**Fig.2.**
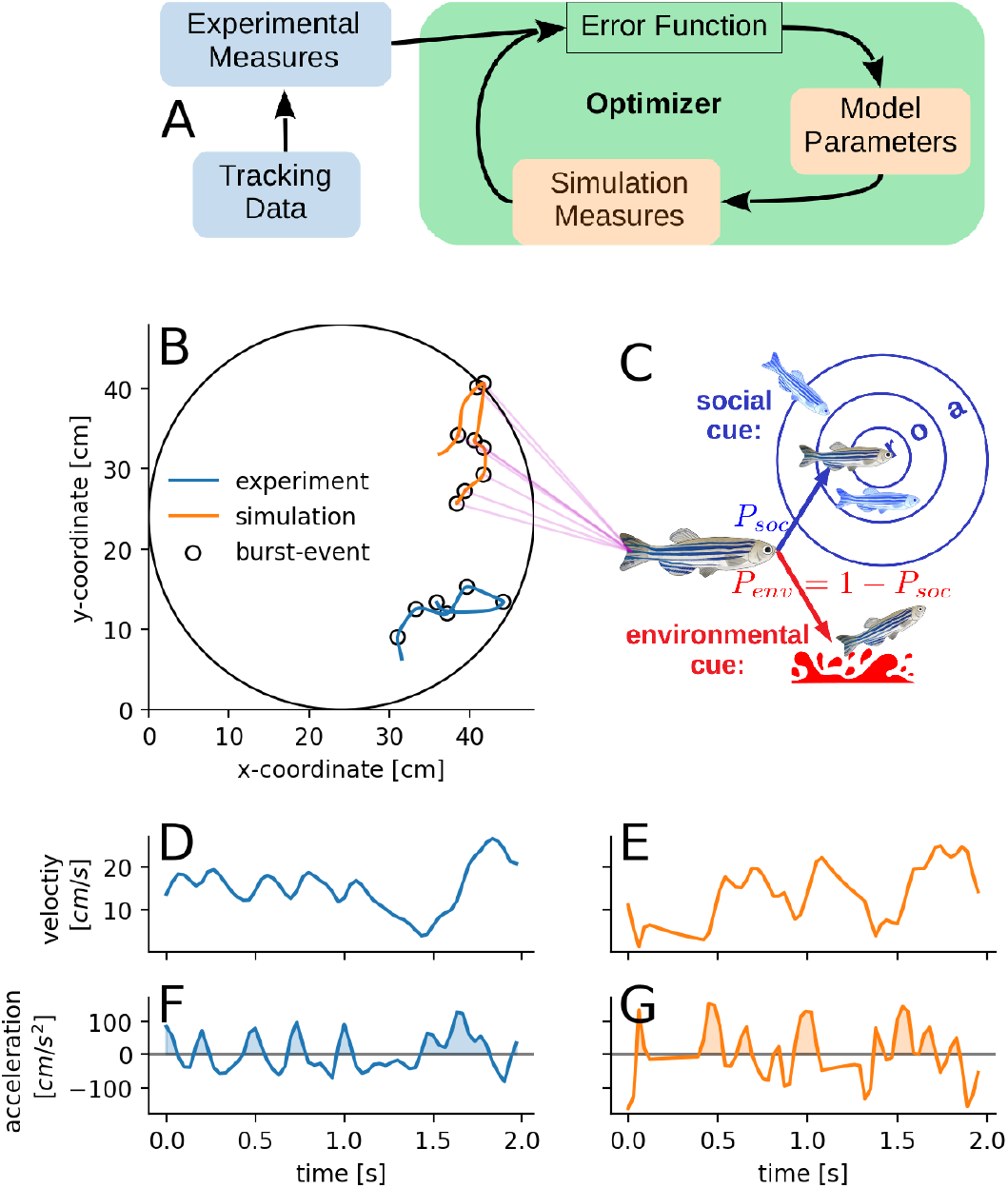
A scheme of the model parameter estimation that was used. The green rectangle represents the optimizer, which updates the model parameters. The update runs until the error function, which compares experimental with simulated measures, is minimal (**A**). The round arena shows representative experimental (blue) and simulation (orange) trajectories of an individual of the random harvested line together with detected burst events (circles; **B**). The schematic model description shows that at each burst an individual can follow with probability *P_soc_* social cues or with probability *P_env_* = 1 – *P_soc_* environmental cues (**C**). The social response of a focal individual to its neighbours depends on how far away they are, i.e. it is repelled (r) from close, aligns its orientation (o) with intermediate and is attracted (a) to neighbours at larger distances, i.e. these zones of behavioral response are exclusive rings around each individual (**C**). The plots show velocity and acceleration of representative experimental (**D, F**) and simulated (**E, G**) trajectories.

**Table 1.** Model parameter overview. The blue colored parameters (burst force, probability of social burst) are estimated by fitting the NND and the mean individual speed of the model simulation to the experimentally observed values. The other parameter could be estimated without explicitly simulating the model. The alignment range *r_o_* of each selection line is identical to the corresponding repulsion range *r_r_*, i.e. there is no alignment zone. The friction coefficient is the same for all selection lines. The lines are abbreviated by LH: large-harvested; RH: random-harvested and SH: small-harvested.

|  | symbol | value |  |  | unit | description |
| --- | --- | --- | --- | --- | --- | --- |
|  |  | LH | RH | SH |  |  |
| repulsion range | $r_r$ | 3.92 | 4.44 | 4.71 | cm | agents closer than $r_r$ repel each other |
| alignment range | $r_o$ | | $r_r$ | | cm | agents at a distance of $r_r \leq r < r_o$ align with each other |
| attraction range | $r_a$ | 15.84 | 18.45 | 19.5 | cm | agents at a distance of $r_o \leq r < r_a$ attract each other |
| burst force | $F$ | 215.1 | 234.4 | 193 | cm/s<sup>2</sup> | acceleration strength during bursts |
| prob. of social burst | $P_{soc}$ | 0.5 | 0.71 | 0.74 | 1 | probability to follow social cues during a burst event |
| burst duration | $t_b$ | 0.089 | 0.091 | 0.097 | s | duration of a burst |
| burst rate | $\gamma$ | 5.7 | 6.3 | 4.7 | 1/s | frequency of burst events per second |
| friction coefficient | $\beta$ | | 2.51 | | 1/s | deceleration strength due to friction (always active) |

We represented the trade-off between vigilance and the use of social cues in the following way. During each burst, the fish decides whether to react to social (i.e., other conspecifics) or random environmental cues with probability *P_social_* and 1 – *P_social_*, respectively. The reaction strength 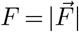 is in both cases the same, only the force direction distinguishes them. The vigilance is the rate of bursts based on environmental information

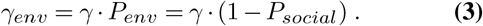

The social force direction 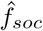 is implemented according to a well-established three-zone model (30). A fish reacts by turning away to avoid collisions at short distances *r* < *r_r_*, by aligning at intermediate distances *r_r_* < *r* < *r_o_*, and by being attracted at large distances *r_o_* < *r* < *r_a_* (Fig. 2C). The direction of the alignment force of fish *i* is the sum of the velocity differences 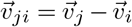 to its nearest neighbors in the alignment zone.

We assume interactions with the nearest neighbors (Voronoi tesselation 51). If multiple zones are occupied including the repulsion zone, alignment- or attraction neighbors are ignored. Otherwise, the force direction is the weighted average of the alignment- and attraction-direction. The weights are the number of neighbors in the respective zone.

The environmental force direction 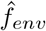 is random if no threat is detected (see Sect. G for possible threats). This is motivated by the fish misinterpreting non-controllable random perturbations (e.g. water reflections) as a threat and responding to them accordingly.

Additional parameters describing the repulsion force from the wall are avoided by a parameter free wall-avoidance mechanism ensuring that the fish avoids the wall while pursuing its intended direction (Movie S2, see SI for details).

An overview with a short description of all model parameters is given in Tab. 1.

#### F. 1. Parameter setting

For the model to resemble the experimental data (Fig. 2B, D-G) we directly measured the parameters if possible, or minimized the error of the simulations with respect to experimental data via an optimizer (Fig. 2A). Six of the eight parameters were inferred from experimental data without the need to explicitly run model-simulation (black colored parameters in Tab. 1). The remaining two (blue colored parameters in Tab. 1) were set by repeatedly simulating the model and reducing the error to the experimental values of the group structure (i.e. NND) and the individual dynamics (i.e. average individual speed), as described below (Fig. S2). We aimed at explaining the differences in NND between the selection lines. Because any of the eight parameters could contribute to the differences, we estimated each parameter for each selection line separately. However, the friction coefficient is assumed to be the same for all lines because it was not behavior dependent and size effects were found to be negligible (see SI Sect. II. A).

##### The simulation-free parameter estimation

was possible for the friction coefficient *β*, burst-rate *γ* and -duration *t_b_* and for the range of each social zone (*r_r_, r_o_, r_a_*). We split individual trajectories in burst and coast phases. A burst phase of a shoaling fish is characterized by an increase in its speed. The friction coefficient *β* is the deceleration during coast phases (Fig. S5). 〈*t_b_*〉 was the mean burst length and 〈*γ*〉 the mean number of burst events per time unit. However, they are different from the model parameters *t_b_* and *γ*. Burst events can overlap each other and consequently reduce the measured burst rate and prolong its duration (Fig. S2D). By a simple binary time series generating model we took account for this effect (See SI Sect. II. A).

We set the range of each social zone by minimizing the angle (error) between the predicted and the experimentally observed direction after a burst (Fig. 2A, SI Sect. II. A). The predicted burst direction of a fish is the social force direction computed from its neighbors relative position and velocity (See SI Sect. II. A).

##### A simulation-based parameter estimation

was necessary for the burst force *F* and the probability to respond to social cues *P_soc_*. We minimized the squared differences (error) between the simulated and experimentally observed values of the nearest-neighbor distances and the average individual speeds. Both measures are emergent properties of the model which require its explicit simulation (Fig. 2A). The differences were standardized by their experimental standard deviation (SI Sect.II. B). We restricted the search space by setting lower and upper bounds for the parameters(see SI Sect. II and Tab. S2). We did not minimize the error to the experimental polarization because (i) it does not differ between selection lines, (ii) the width of the orientation zone, mostly influencing the polarization, is already estimated from data, (iii) boundary effects could affect it (i.e., fish might be attracted to the wall but avoid it at close ranges, which could lead to a confounding alignment with the wall). For the minimization we applied the covariance matrix adaptation evolution strategy (CMA-ES; (52)), from the Python package *pycma* (53) which is a good choice for a multi-modal, noisy error-landscape.

### G. Predator and fishing simulations

We used the agent-based model representations of the three selection lines to investigate whether the size-selective harvest could impact the ability of the shoals to evade a natural predator and different fishing gear. We simulated *N* = 30 shoaling fish in a box of size *L* = 100 *cm* with periodic boundary conditions. Three scenarios were considered: (i) natural predation - including the confusion effect (14) - by a single mobile predator following the closest fish; (ii) fishing by a single agent without information about the position of the shoal (random search, similar to a fisher on a boat without an echo sounder); (iii) fishing by multiple aligned agents moving on a straight line towards the center of mass of the shoal (e.g., commercial trawling informed on the position of the shoal).

In contrast to the simulation with only shoaling fish, now the environmental force direction 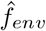 is given by a direct avoidance force (repulsion) away from the simulated predator/fishing agent, if it is detected. The detection probability is maximal up to the detection distance *r_f_* = 7 *cm* ≈ 3*BL*, and it decays linearly until it equals zero at *r* = 35 *cm* ≈ 14*BL*.

All additional parameters for the three scenarios are explained below (summarized in Tab. S3).

#### G.1. Natural predator

The predator moves directly to the closest fish (Movie S3) with *v_net_* = 20, 25, 30, 35 *cm/s*, which is larger than the average speed of shoaling fish (〈*v*〉 ≈ 15 *cm/s*) because predators are usually larger than prey and therefore can swim faster (54, 55). Most predators attack a specific fish and therefore need to focus on it before attacking. The so-called confusion effect (56), the disruption of the predator focus by a large number of individuals who are difficult to distinguish by phenotype and movement, is believed to be one key benefit of group living (12). We model the probability of a prey to be successfully captured if it is closer to the predator than *r_capture_* = 5*cm* within a small time window [*t,t* + *δt*] as

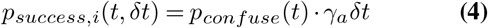

Here *γ_a_* = 1/*s* is a base predator attack rate and *p_confuse_* (*t*) represents the confusion effect that modulates the attack rate. The confusion term decreases *p_success_* with increasing number of perceived prey *N_sensed_* in a sigmoidal fashion (see SI sect.I. D). Thereby, *N_conf_* =4 is the number of sensed shoaling fish at which *p_confuse_* = 0.5 (56), and a fish is sensed if it is closer than *r_sense_* = 4*r_capture_*.

#### G.2. Fishing agents

Fishing agents always capture a shoaling fish if it is closer than *r_capture_* = 5 *cm*. Fishing gears were simulated by varying the speed of the fishing agents (*v_net_* = 7.5, 15, 22.5 *cm/s*).

In the single fishing agent scenario (Movie S4), the agent performs a random search with constant speed *v_net_*. Its angular change is given by 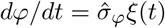 with *ξ*(*t*) as white noise. For different speed values the angular noise strength 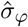 is adapted such that the persistence length is constant 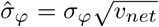, i.e. the change in angle after travelling a certain distance is independent of its velocity (see SI sect.I. E).

In the multiple aligned fishing agents scenario (Movie S5), the agents (*N_f_* = 50; red) are aligned on a line spanning *L/*4. After travelling for a distance *L/*2, the fishing agent array is recreated at a distance of *L*/4 away from the center of mass of the shoal and restarts its movement in its direction.

## Results

### A. Shoaling behavior

We found that mean inter-individual distance was repeatable across time (*R_adj_* = 0.51[0.21 – 0.79]; *p* < 0.001; group tracking at 150-190 days post fertilization). The results of the individual tracking at 190 days post fertilization indicated a significant effect of the selection lines on mean inter-individual distance, independently of measuring it in cm (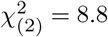; *p* < 0.05) or mean body length 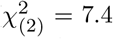; *p* < 0.05; Fig. 3A). By contrast, we found a trend of selection lines on mean nearest-neighbor distance when measuring it in cm 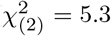; *p* = 0.070; Fig. 3A), while it had a significant effect when measuring it in body length 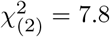; *p* < 0.05; Fig. 3A). In particular, the large- and small-harvested lines tend to form less and more cohesive shoals than controls, respectively (Fig. 3A). Moreover, we did not find a significant effect of selection lines on group polarization (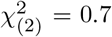; *p* = 0.716; mean and standard deviation across selection lines: 0.40 ± 0.06). We also found a negative correlation (*Rc* = –0.38; *F*_(1,28)_ = 12.5; *p* < 0.01) between mean nearest-neighbor distance and the previously documented group risk-taking behavior (Fig. 3B; note that in this study we used the same groups of fish previously used to test group risk-taking behavior (7), but tested at different days post fertilization). Furthermore, the inter-individual distance and the nearest-neighbor distance also showed a strong positive correlation (*Rc* = 0.88; *F*_(1,28)_ = 207.2; *p* < 0.001), which means that the change in inter-individual distance was not related to a split of the main group into multiple subgroups.

**Fig. 3.**
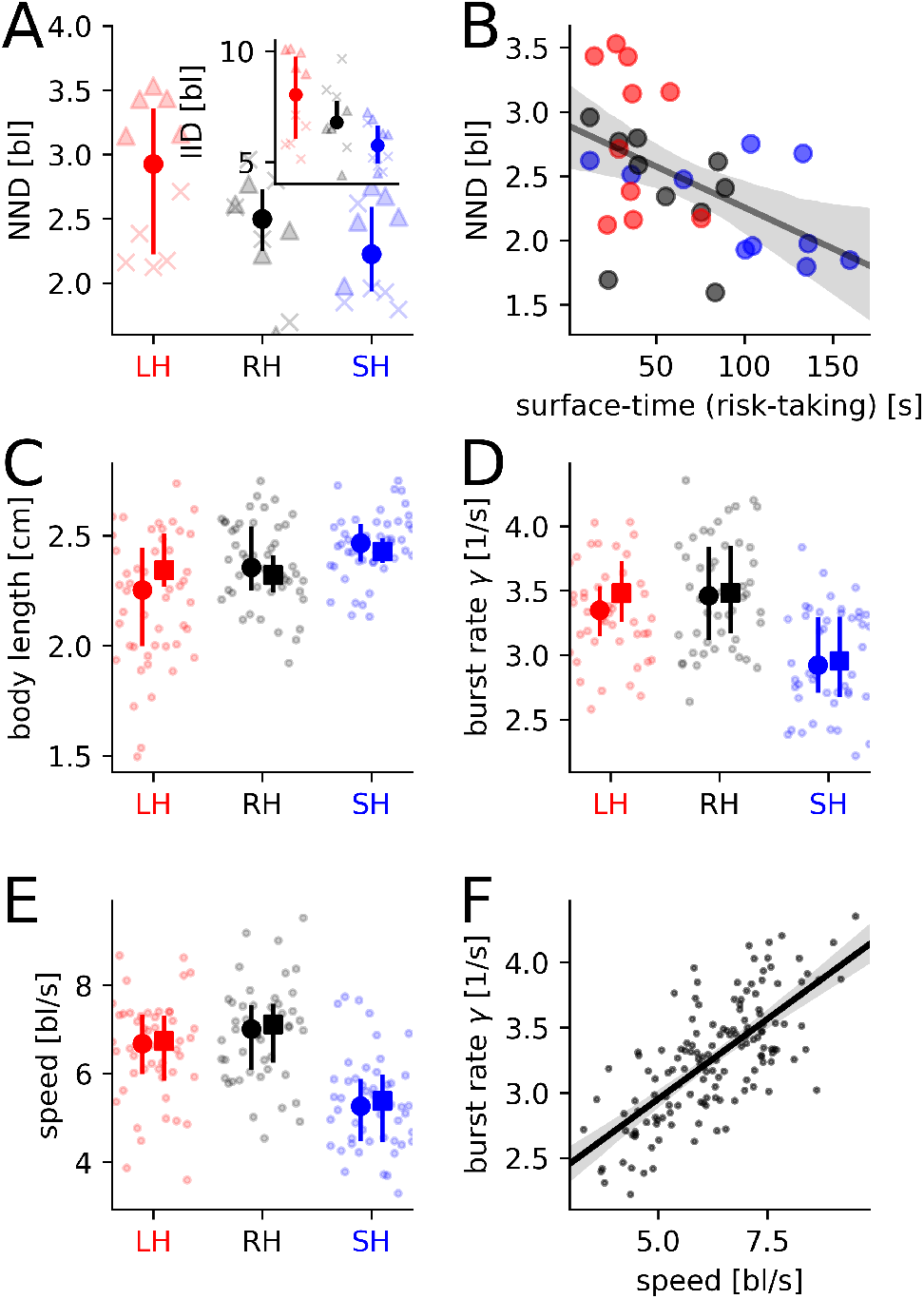
Group level results at 190 days post fertilization for mean nearest-neighbor distance (in body length; **A**) and inter-individual distance (in body length; inset **A**) together with the correlation between mean nearest-neighbor distance and previously documented group risk-taking behavior (i.e. time spent feeding at the surface at 230-240 days post fertilization, **B**; see (7) for more details). Individual level results for body size (**C**), burst rate (**D**), and speed (expressed in body length per second; **E**) as well as the correlation between burst rate and speed (**F**). The selection lines are indicated by different colors (red=LH: large-harvested; black=RH: random-harvested; blue=SH: small-harvested). Vertical lines with circles (**C-E**) refer to the entire data set (N = 48), while lines with squares refer to the subsample of size-matched individuals (N = LH: 24; RH: 36; SH: 40). The bold circle and square represent the median and vertical lines represent the first (25th percentile) and third (75th percentile) quartiles. Triangles and crosses (**A**) correspond to raw data of the two different replicates.

### B. Fish body size and individual behavior in the shoal

As expected, the body size at age of the experimental fish varied significantly among the selection lines (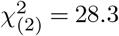; *p* < 0.001; Fig. 3C). The individual burst rate within the shoal was significantly (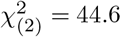; *p* < 0.001) different among selection lines. Specifically, the small-, but not the large-harvested line, bursted less frequently than controls (Fig. 3D). Furthermore, the individual swimming speed was significantly (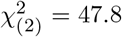; *p* < 0.001) different among selection lines. Again, the small-, but not the large-harvested line, showed a slower swimming speed than controls (Fig. 3E). Note that the burst rate showed a positive correlation with the swimming speed (*R* = 0.51; *F*_(1,142)_ = 148.6; *p* < 0.001; Fig. 3F), which suggests that differences in burst rate was the main cause of variation in swimming speed.

To control for possible confounding effects of body size, we composed a subsample of the dataset with size-matched individuals without significant (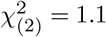; *p* = 0.592) differences in body length among the lines (Fig. 3C). The above mentioned differences in speed and burst rate among the selection lines were still significant for both burst rate (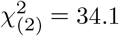; *p* < 0.001; Fig. 3D) and speed (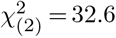; *p* < 0.001; Fig. 3E) when the subsample of size-matched individuals was used in the analyses. Moreover, body size did not relate to burst rate (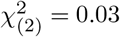; *p* = 0.858) and swimming speed (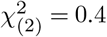; *p* = 0.519).

### C. Mechanistically linking micro-level interactions among individuals with macro-level collective outcomes

We allowed each parameter except of the friction coefficient to vary between the model representations of the selection lines (parameters summarized in Tab. 1). The selection lines were about equal in burst duration (*t_b_*) and the differences in burst rate (*γ*) corresponded qualitatively to the experimental equivalents (Fig. 3D). The range of the social zones (*r_r_, r_o_, r_a_*) differed in the same manner as the experimentally observed body lengths (Fig. 3C), i.e. the on average smaller fish (LH line) have a shorter repulsion, orientation and attraction zone and vice verse for the larger fish (SH line). The probability to follow environmental cues (*P_env_*, Fig. 4A) and the burst force (*F*, inset Fig. 4A) were set such that the model representations of nearest neighbor distance (Fig. 4C) and mean individual speed (Fig. 4D) were as close as possible to the experimental data. Interestingly, the closely related vigilance (product of burst rate and *P_env_*) was greater than control for the large-harvested line, and lower for the small-harvested line (Fig. 4B). The same qualitative differences were reported for group risk-taking behavior in a previous study, where behavior was measured at different days post fertilization ((7);, see also Fig. 3), thus results are robust. A possible mechanistic explanation for the observed differences in cohesion is that the bolder individuals of the small-harvested line were less vigilant, which causes them to respond less frequently to environmental cues, in contrast to social cues, which consequently increased their cohesion (vice versa for the large-harvested line). To substantiate this explanation we re-estimated the parameters *F, P_env_* by enforcing that all selection line shared the same *P_env_*. Without the ability to differ in *P_env_*, and thus in vigilance, the model representations were unable to reproduce the experimentally observed cohesion pattern (Fig. S12).

**Fig.4.**
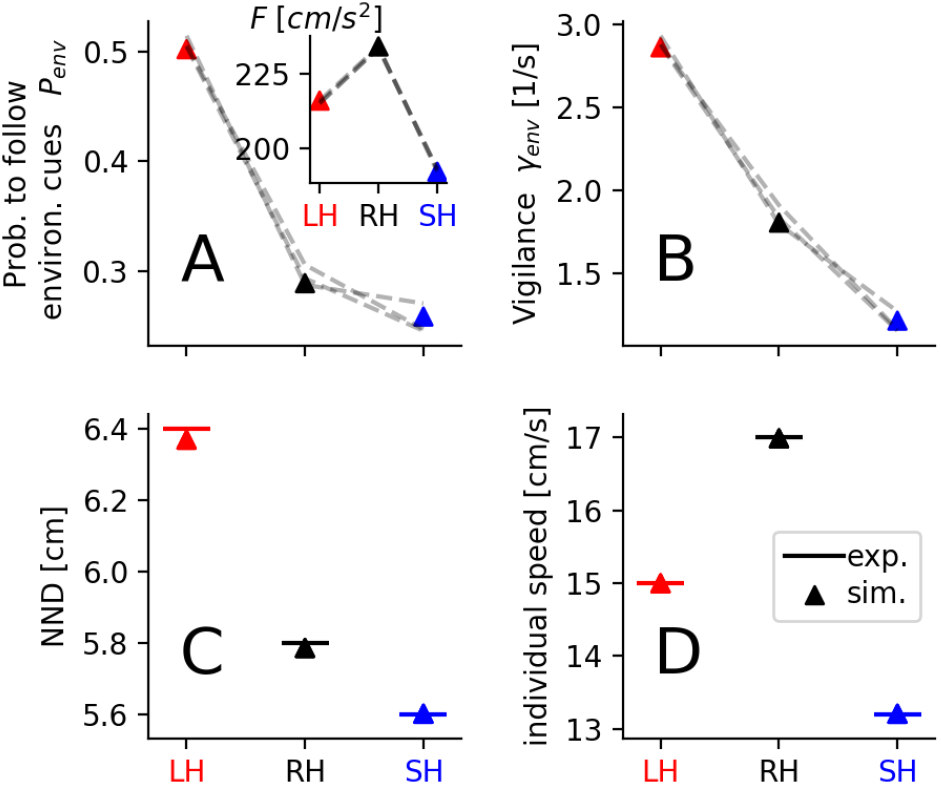
Fitted model representation of the different selection lines. In order to resemble the experimental observations the probability to follow environmental cues (**A**) and the burst force (inset **A**) were set for each selection line. The vigilance *γ_env_* is the product of *P_env_* and the burst rate *γ* (**B**). The average nearest neighbor distance (NND, **C**) and the average individual speed (**D**) are emergent properties of the model and were used to quantify how well the parameters (*P_env,γ_*) reproduce the experimental observations. Triangles represent the model parameters or the simulation outcomes of the parameter set with the best match to the experiment. Dashed lines (**A, B**) represent a parameter set of a different initialization, and therefore show the robustness of the best matching parameter set (triangles). The horizontal solid lines (**C, D**) represent the experimental values. Note that in contrast to Fig. 3 units are in *cm* because the modelled agents have no body length. The selection lines are indicated by different colors (red=LH: large-harvested; black=RH: random-harvested; blue=SH: small-harvested).

### D. Mortality rates in the predator and fishing contexts

The simulation results showed that if the natural predator moved slow, both size-selected lines had a higher mortality rate than controls (Fig. 5C). By contrast, when the predator moved faster, the large-harvested line experienced higher mortality than the controls, while the small-harvested line did not showed major differences. In both fishing scenarios (Fig. 5D, E), the large-harvested had a lower mortality rate than the control, and vice versa for the small-harvested line (Fig. 5D, E). Note that when the confusion effect was turned off, the mortality rates observed in the natural predator scenarios were similar to those of the fishing scenarios (Fig. S3B). It suggests that the differences in group cohesion among the selection lines (Fig. 4C) influenced the mortality rate of the natural predator (i.e., the confusion effect is stronger in more cohesive shoals because individual fish are at lower risk in denser regions, which could be applied also to relative position within the shoal). The confusion effect as explanation is further supported by the average distance of the shoaling fish from the predator/fishing agent. The distance was greater than control in all the scenarios for the large harvested line, and vice versa for the small harvested line (Fig. S3 I-P). Thus, the fish shoals avoid the natural predator as well as they avoid the fishing agents but the predator is confused by the cohesive shoaling (for details see SI Sect. III).

**Fig. 5.**
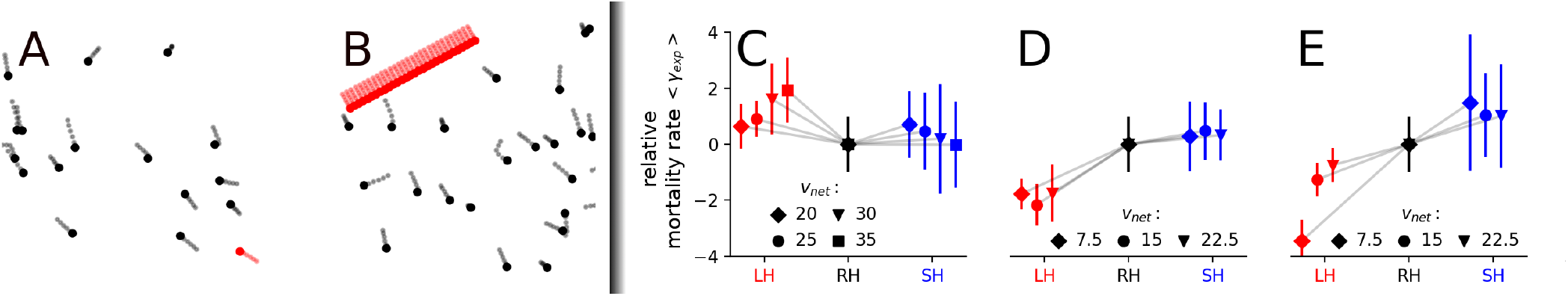
Simulation snapshots are shown for the predator- and single-fishing-agent scenario (**A**) and for the multiple-fishing-agents scenario (**B**); with shoaling fish in black and the predator and fishing agents in red. The mortality rate, i.e. prey captured or fished per time unit, was computed for a natural predator moving with *v_net_* = [20, 25,, 30, 35] *cm/s* (**C**); for a single fishing agent (**D**) and for multiple fishing agents distributed on a line (**E**). The fishing agents moved with *v_net_* = [7.5, 15, 22.5] *cm/s* indicated by rhombus, circle and triangle, respectively. The shoaling fish had a mean speed of about 〈*v*〉 ≈ 15 *cm/s*. The markers (**C, D, E**) represent the relative mortality rates, i.e. the mortality rates were reduced by the mean and divided by the standard deviation of the random-harvested line. The markers and error bars indicate the mean and standard deviation estimated from samples of 400 simulations of *N* = 30 shoaling fish in a box of size *L* = 100 *cm* with periodic boundary conditions. The colors red, black and blue correspond to the large- (LH), random- (RH) and small-harvested (SH) selection line, respectively.

## Discussion

We showed that five generations of size-selective harvesting followed by eight generations during which harvesting halted left a legacy in terms of shoaling behavior. Shoal cohesion changed in the direction expected by the trade-off between vigilance and the use of social cues (12). In particular, using the agent-based model, we revealed a decrease of vigilance in the small-harvested line that was linked to an increase of attention to social cues, leading to more cohesive shoals (vice versa for the large-harvested line). We also explored possible consequences for fishing and natural mortality. Specifically, the shoaling behavioral changes induced by large size-selective harvesting (i.e., a typical selectivity pattern in many fisheries) may decrease mortality in fishing scenarios, but increase natural mortality.

The results of the burst-coast, agent-based model linked boldness with shoal cohesion and provided a mechanistic explanation of harvesting-induced changes of shoaling behavior by assuming the existence of a trade-off between social and environmental information in collective motion (57). Our model is similar to previous work in terms of splitting the movement of shoaling fish into an active and passive phase (27, 39, 58). However, it provides a novel perspective about collective movements of fish shoals by linking risk perception and movement decisions. In fact, most previous models of collective behavior only accounted for social information (27, 30, 58–60), while our model explicitly implements the trade-off between random environmental and social cues. The previous simulation studies which considered similar mechanisms (e.g., when moving agents directly react to a non-conspecific cue as a predator (61, 62) or a food-patch (63)) allowed the moving agents permanent access to non-social information without any detriment to the perception of social cues, i.e. without any attention trade-offs. In our model, the vigilance limits the general capacity to react to social cues and thus affects the individual behavior even in the absence of environmental cues. Another major difference to most existing models (27, 30, 60, 64) is that the individual speed is not a model parameter but emerges from the interplay of rate, strength and duration of bursts with the tendency to follow environmental cues. Especially, accounting for the possibility of individual burst being triggered by either social or environmental cues adds a novel, yet ecologically relevant behavioral dimension to our agent-based model.

Our integrative research approach focused on the impact of size-selective harvesting on individual fish behavior and consequently on the emergent collective behavior. We did not considered changes of instantaneous shoaling behavior mediated by a direct link between perceived risk and shoal cohesion (i.e., the higher the risk the more cohesive are the shoals; (37, 65)). In that context, an increase of boldness in the small-harvested line could have been linked to a decrease of perceived risk and consequently to a decrease of shoal cohesion. However, our experimental results showed the opposite pattern reinforcing the idea that size-selective harvesting can have an evolutionary impact on shoaling behavior mediated by a trade-off between vigilance and the use of social cues. The changes of shoal cohesion we documented may also be interpreted in light of a behavioral-mediated resource acquisition mechanism that affected size-at-harvest as trait under selection (7, 66). Zebrafish can attain a higher food consumption rate when they are in small groups than alone (38). Therefore, the large size-selective harvesting treatment could have removed individuals that fed in small groups because such behavior could have provided advantages in accessing food and hence increased size-at-harvest; this could have fostered the evolution of reduced shoal cohesion (and vice versa for the small size-selective harvesting). However, previous results on the selection lines showed that when assaying individual female sociability, the small-harvested line was found to be less social than control, while the large-harvested line was not found to differ from controls (42). Thus, we suggest that direct selection on sociability was not a major driver of the changes in shoal cohesion documented here.

The simulations of natural predation and fishing revealed that the shoaling behavioral changes of the large-harvested line could affect both fishing and natural mortality. Specifically, in our simulations the increase in vigilance of the large-harvested line decreased vulnerability of shoals under a multitude of simulated fishing scenarios considering single and multiple fishing agents moving at different speed. Fisheries-induced evolution of group behavioral traits (i.e. shoal cohesion) can have strong repercussions in many shoaling teleosts when attempting to reduce risk of natural predation (15). Our study suggests that intensive harvesting directed at the larger individuals of a population could shift the fitness optimum of shoaling behavior in the opposite direction of what natural selection would favor. Evolution under anthropogenic selection has been demonstrated to occur within few generations (3–5, 67–69), and could impede recovery of exploited population even after harvesting halted (70, 71). Here, we provide the first functional integration of individual vigilance into mechanisms governing group dynamics with respect to fisheries and natural predation.

An alternative mechanism, recently proposed by Guerra et al. (9), is purely based on group size, which is expected to decrease in response to fishing harvesting and subsequently decrease fitness in response to natural predation. Therefore, it is plausible that fisheries-induced evolution affect multiple traits related to shoaling behavior that could vary depending from the specific fishing gear used and the ecology of the exploited species (72). Our work proposes a new mechanism that could increase the natural mortality associated with fisheries-induced evolution and thereby negatively affect recovery, even if harvesting halted.

Previous studies have shown that fisheries-induced evolution of life histories can substantially increase natural mortality (73, 74). Elevated and size-selective harvesting promotes the evolution of a fast life-history, characterized by early maturation and small size, elevated reproductive investment and reduced post-maturation growth (75). Natural mortality decreases with length in fishes (2); therefore a decrease in body size is linked to an increase in natural mortality. In addition, an increase in reproductive investment is associated to an increase in natural mortality due to the cost of spawning (73, 76). Moreover, elevated and unselective fishing harvesting is expected to increase boldness, which in turn increases natural mortality (73). Previous experimental results using the zebrafish selection lines indicated that elevated mortality in combination with large size-selective harvest decreased group risk-taking behavior (7), which agrees with recent theoretical work (6). Yet, while decreased boldness might reduce natural mortality, we propose that the decrease in shoal cohesion emerging from the evolution of vigilance can be a further mechanism by which natural mortality increases in a fisheries-induced evolution context where large individuals are preferentially harvested. We therefore extended the framework presented by Jørgensen and Holt (73) for unselective fisheries harvest revealing an insofar unknown mechanism by which fisheries-induced evolution of individual behavioral trait through size-selection may drive collective outcomes that may affect natural mortality in shoaling fishes. This is particularly important in an ecosystem context where trophic flow between predator and prey is controlled by a bottom-up mechanism mediated by prey behavior (77).

The interpretation of our results has limitations. First, the selection lines were not exposed to predation pressure during the selection experiment. The evolutionary trajectory of shoaling behavior could have been dampened by the presence of predation pressure and could reduce the effects described here in particular for the large-harvested line. Second, shoaling behavior is inherently complex and other factors that were not considered here could play a major role. For example, time spent feeding could be traded-off with risk of predation (15, 16), or shoal cohesion could be directly affected by light levels (17). Most importantly, we cannot exclude that presenting a real predation risk during the shoaling trial could have changed our results. Such aspect is now under study in a follow up experiment with the zebrafish selection lines. More research that extends to other mechanisms or ecological conditions is warranted to test the robustness of our findings. Third, we measured the impact of size-selective mortality at the phenotypic level without providing genetic support for evolution. However, previous analyses revealed that genetic changes have indeed taken place in the zebrafish selection lines (5), and we also demonstrated here that shoaling behavior was a repeatable trait, which suggests that it is potentially heritable. Despite such limitations and the fact that we focused on a model species that mainly shoals in small groups, our study expanded a recently published framework on the effects of fishing on obligate schooling fish (9). We used a non-obligated schooling species (34), which is a behavior that may convey adaptive value in the wild (35, 36), and also showed a response to size-selective harvesting over multiple generations. This could render our results of a wider interest for fisheries where many important exploited species are non-obligated schooling species (78).

## Conclusions

There is an ongoing debate on the role of fisheries-induced evolution on population dynamics (79–81). Our results suggest that fisheries-induced evolution of shoaling behavior could increase natural mortality and therefore could indirectly contribute to slow down recovery of exploited population in terms of population biomass. Our results provide an experimental baseline for understanding the impact of size-selective harvesting, as well as other similar patters where survival is strongly influenced by body size, on shoaling behavior. We hope our study stimulates more research on the effects of fishing and other stressors on shoaling behavior and possible consequences for population dynamics and fisheries.

## Data and codes availability

Experimental data are available as supplementary material together with the R code used for statistical analysis. The code to run the burst and coast model with the fishing/predator scenarios is available at github (https://github.com/PaPeK/sde_burst_coast). The optimization was done with a python implementation of the CMA-ES (https://github.com/CMA-ES/pycma).

## Contributions

V.S. and R.A. designed the experiments; V.S. performed the experiments, the statistical test and analysed part of the experimental data. P.P.K. and P.R. developed the model. P.P.K. ran the simulations and analyzed part of the experimental data. V.S., P.P.K., P.R. and R.A. interpreted the data. V.S., P.P.K, P.R. and R.A. wrote the manuscript.

## SI Appendix Supplementary Information

### I Model description

The burst-coast model intends to mimic the burst-coast swimming behavior of zebrafish (*Danio rerio*). We assume that a fish is accelerating only during the burst phase with a constant force of magnitude *f_b_*, while no forces are present during the coast phase implying a deceleration due to friction. The differential equations of motion defining current position 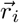 and velocity 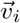 of an agent *i* are:

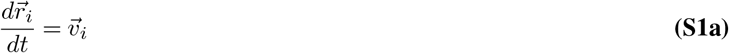

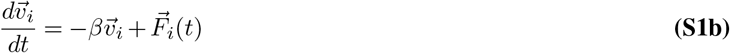

with 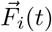 as a finite social or environmental force vector with |*F*(*t*)| > 0, for fish in the bursting phase. A fish decelerates passively during the coasting phase, thus the force vector vanishes 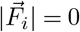. *β* is the friction coefficient. The velocity change of Eq.(S1) can be split into the part parallel and perpendicular to the current velocity direction 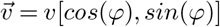:

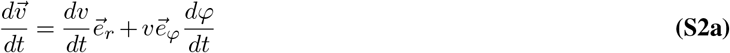

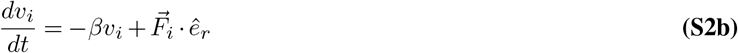

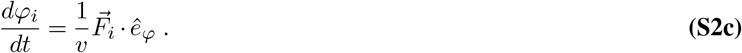

Note that the *v*^−1^ dependence of the turning rate (Eq. S2c) follows directly from Eq. S1b via a coordinate transformation (50). In addition it has been verified by experimental tracking data (Fig. S4).

The burst behavior is defined by the burst rate *γ*, the burst duration *t_b_* and the burst force 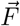. In particular the burst force 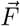 governs whether the fish uses social or environmental cues. The fish decides at the start of a burst with a probability *P_social_* whether to react socially to other fish or, with probability 1 – *P_social_*, to environmental cues.

The resulting burst force 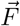 is

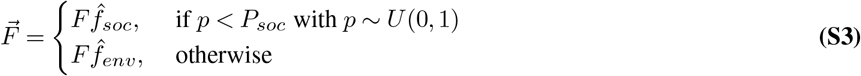

with *F* as the magnitude of the burst force and 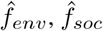 as the unit vectors in direction of the environmental or social cue.

#### I. A. Social forces

The social force is motivated by a three zone model (30) consisting of a repulsion zone vanishing at a distance *r_r_*, followed by the alignment zone vanishing at *r_o_*, and the attraction zone vanishing at *r_a_*. Thus, *r_r_ < r_o_ < r_a_*. The direction of the social forces is computed by

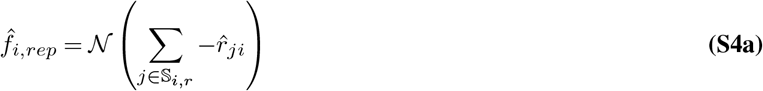

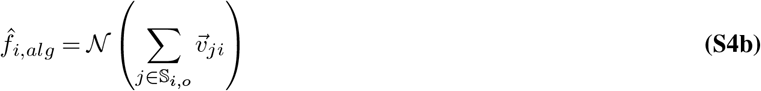

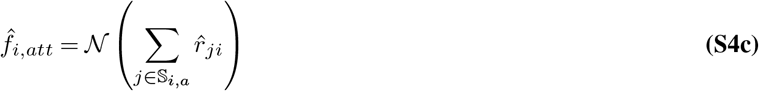

with 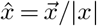 defining a unit vector, 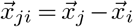 as the difference between the vectors of fish *j* and *i*, 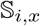 is the set of indices of fish in zone *x* of fish *i*, and 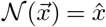 is a normalization operator. We assume Voronoi interactions because they provide a reasonable approximation of visual networks (51) and can be efficiently computed. Therefore, the sets 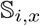 with *x* ∈ [*r, o, a*] are composed only of Voronoi neighbors of fish *i*. Note that the alignment force is the sum of the velocity difference vectors 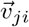. Thus, the focal fish *i* experiences the strongest alignment with neighbours whose velocity vectors differ the most from its own. If neighbors of fish *i* occupy different zones simultaneously the following rules apply:

- if 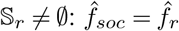 (repulsion dominated)
- 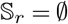 and 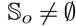 and 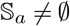: 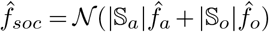 (weighted average)

#### I. B. Environmental force

In the absence of a predator the environmental force is modelled as a random force vector. This models the sensitivity of individual fish to environmental noise (e.g. water reflections, water perturbations, sounds), inducing false-positive escape responses. In the presence of a threatening agent the environmental force is modelled as a simple repulsion of the shoaling fish from the simulated predator or fishing agents

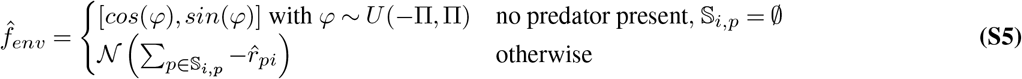

with *U*(*a, b*) being a uniform distribution with *a* and *b* as lower and upper bounds, and 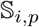 being the set of predator or fishing agents detected by a shoaling fish *i*. A predator or fishing agent is detected by a shoaling fish with probability

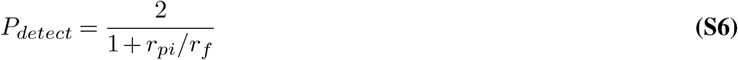

which equals one at *r_pi_* = *r_f_*. If *P_detect_* > 1 it is set to 1. *r_f_* is the detection distance which was for all simulations *r_f_* = 7 *cm*. This value is reasonable because fish should be able to detect a predator when they are likely to be captured *r_f_* ≥ *r_capture_* = 5 *cm* but the distance should also be close to *r_capture_*, otherwise fish would respond too often to non-dangerous cues and because the visibility in water decays with distance. The randomness is a consequence of the assumption that shoaling fish misinterpret reflections on the water surface as threats since there were no “real” threats in the experiment.

#### I. C. Wall-avoidance mechanism

We attempt to design the model as close to the experimental setting as possible and therefore use a circular boundary. The introduction of parameters describing the repulsion force from the wall was avoided by the fish following a parameter free wall-avoidance mechanism. It is based on predicting before each burst the future position 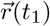 of the shoaling fish at the next burst (at *t_t_nb* which stands for “time to next burst”) plus some extra time (at time *t*_1_ = *t_tnb_* + *t_b_*). The extra time *t_b_* is necessary because the agent can not prevent collision with the finite burst force, if it is at the next burst inside the tank but directly at, or very close to the wall. The length of the extra time is set to the burst time *t_b_*, because if the agent coasts (i.e. uses no force) and does not collide with the wall, any burst force is sufficient to prevent collision if the agents bursts instead. If the predicted position 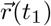 is outside of the tank, the force direction is adapted until 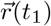 is inside the tank. Thereby is the smallest possible change in force direction used. This ensures that the shoaling fish avoids the wall while pursuing its intended direction.

To predict the future position 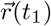 with *t*_1_ = *t_t_nb* + *t_b_* the solution of the coupled differential Eq. S1 for 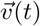 and 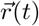 are used. Since the x and y components of 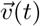 and 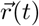 are not coupled with each other, the solutions are computed for each component separately:

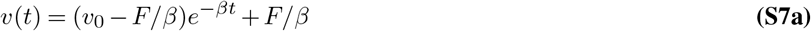

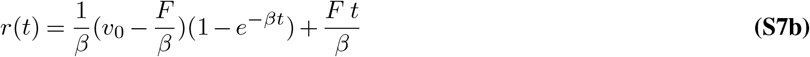

where we omitted the index *r_x_* or *r_y_* for the terms *r, F, v*_0_ and *v* for simplicity. The solutions above assume a constant force *F*. Therefore, first, the position and velocity after the burst are predicted and then, second, the position after the coast.

#### I. D. Capture probability of a single fish by a natural predator

If a shoaling fish is closer to the predator than *r_capture_* its probability to be successfully captured within a small time window [*t, t* + *δt]* is

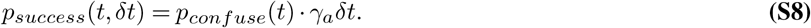

Here *γ_a_* is the predator attack rate, and *p_confuse_* (*t*) represents the confusion effect that modulates the attack rate. The attack rate used in the simulations is *γ_a_* = 1/*s*. The confusion term depends on the spatial configuration of shoaling fish around the predator and decreases *p_success_* the more individuals are sensed (14). A shoaling fish is sensed if it is closer than *r_sense_* = 4 · *r_capture_*. We assume that it modulates the successful capture probability in a sigmoidal way depending on the number of sensed shoaling fish *N_sensed_*:

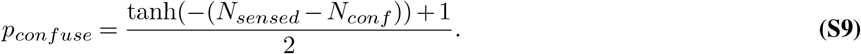

Here *N_conf_* = 4 is the number of sensed shoaling fish at which *p_confuse_* = 0.5.

#### I. E. Random movement of angling agent with constant persistence length

The individual fishing agents perform stochastic movement, with a fixed persistent length independent on their speed. The evolution of the heading angle is given by

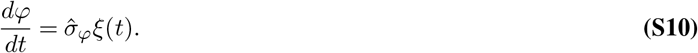

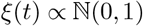 is Gaussian white noise with zero mean and variance of one. 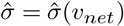 is the angular noise strength whose exact form and dependence is derived below.

Keeping the persistence length constant is equivalent with keeping the variance in angle constant after the individual fishing agent travelled a path of length = *l*. The time needed to travel this path-length is *t_l_* = *l/v_f_* and the variance in angle after travelling the path is

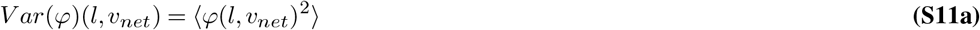

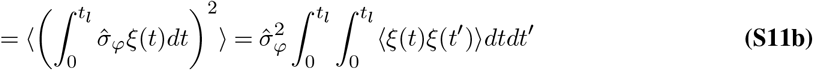

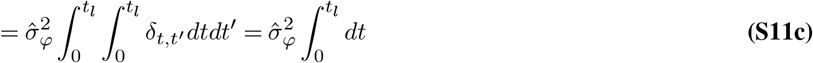

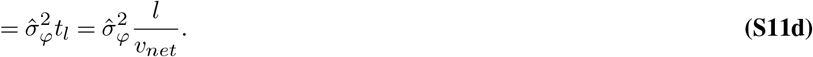

Here, we used uncorrelated Gaussian white noise, i.e. 〈*ξ*(*t*)*ξ*(*t*′)〉 = *δ_t,t′_*, with *δ_t,t′_* as the Kronecker-delta. If we set 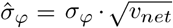, the variance in angle after travelling a path of length *l* is independent of the speed:

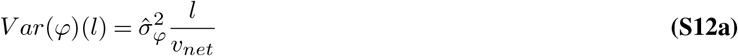

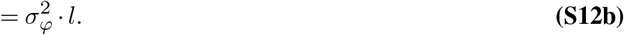

### II Parameter setting

The model has in total eight parameter, from which six can be directly estimated from experimental measures without the need to simulate the model. The remaining two parameters can not be directly accessed from data and were estimated by an optimization. The optimization minimized the sum of squared differences between model emergent properties and their experimental measured values. For an overview of all model-parameters and how they are set see Fig. S2A. First, section II. A explains the simulation-free parameter estimation from experimental measures in detail. Second, section II. B explains the optimization of the remaining parameters by comparing model emergent measures. All model parameters are summarized in Tab. 1.

#### II. A. Simulation-free parameter estimation

In the model (see section I) an agent can be either in the burst or coast phase, thus we decompose individual trajectories into these phases. Burst and coast phases are characterized by an increase and decrease in the speed of the shoaling fish, respectively. In the coast phase the only parameter which defines change in velocity is the **friction coefficient** *β* (Eq. S2). Thus, we estimated *β* = 2.51 by the slope of the linear regression between the individual speed and the average deceleration during coast-phases as shown in Fig. S5. Because the friction force is by definition zero at zero velocity we fixed the interception point to zero and therefore found effectively the least-square solution of the linear equation 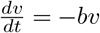. We assumed the same friction coefficient *β* for the model representations of all selection lines because it is reasonable to assume that friction properties will primarily depend on the cross-section of the fish perpendicular to its their moving direction. The size differences between selection lines could in principle change the friction (larger individuals have a larger cross section and therefore a larger friction coefficient), but the differences probably negligible. Note, that we also assume the same mass for all selection lines and therefore could different average masses also influence the effective friction (which is strictly the friction divided by the assumed mass). Second, even if they are not negligible or differ in mass, a lower friction coefficient does impact the speed in the model as does the burst-force. Thus, by allowing the burst force to vary between selection lines we effectively take account for differences in the friction coefficient.

The average **burst duration** 〈*t_b_*〉 and **rate** 〈*γ*〉 were estimated by the mean length and frequency of burst periods. For the large-, random- and small-harvested line the burst duration was 〈*t_b_*〉 = [0.117,0.123,0.123]*s* and the burst rate was 〈*γ*〉 = [0.35,0.44,0.38], respectively. In the model a new burst period can start during an already ongoing burst and therefore prolong the measured burst duration and decrease the rate, as illustrated in Fig. S2D. In consequence, the average estimated burst duration and rate are different from the model-parameters *t_b_* and *γ*. We approximated them by creating a binary time series in which at each time-step *dt* a burst-event happens with probability *γ* · *dt* which raises the acceleration from zero to *F* for a duration of *t_b_*. We simulated this process, computed the average values 〈*t_b_*〈*N*,〉 〉*γ*〈*N* (as seen in Fig. S2D) and selected the *γ* and *t_b_* which minimized the summed square differences to the experimental observed averages for each harvested line. In Fig. S1 the squared differences are shown. The resulting parameters are reported in Tab. 1.

The **burst force** *F* in heading direction *f_b,s_* can be estimated by adding to the change in speed 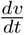 the friction coefficient times the current velocity *β* · *v*. This results in the mean burst force in heading direction *f_b, s_* = 95*cm/s*^2^. However, from Eq. S2 it is clear that the total magnitude of the burst force is defined as

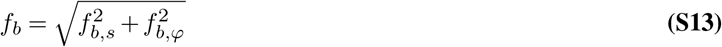

with 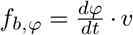 (from Eq. S2c). Note, that we are using *f_b_* instead of the symbol for the model parameter *F* because we expect *F* to be larger than *f_b_* since the latter is estimated from forces only during the burst-phase. However, the turning force often started prior to the acceleration in velocity direction and reached its maximum at the start of the burst (Fig. S6C, D). This suggests that the actual force *F* needed to mimic the characteristic zigzag-like swimming of zebrafish is larger than *f_b_*. For the different selection lines we estimated 〈*f_b,LH_*〉 = 121.2*cm/s*^2^, 〈*f_b,RH_*〉 = 135.2*cm/s*^2^, 〈*f_b,SH_*〉 = 102.3*cm/s*^2^. Thus, instead of setting *F* = *f_b_* we used its largest mean to set the boundaries of the search-space for *F* in the optimizer.

The **ranges of the social-interactions zones** (*r_r_*, *r_o_, r_a_*) are set to minimize the angle between the predicted and the actual direction after a burst (see Fig.S2C). The computation of the predicted direction is based on the relative positions of neighbors of the focal, bursting fish at the start of the burst. From this neighbor constellation the direction of the social force is computed with Eq.S4a. This is done for each burst-event in which the bursting fish was at least three body length away from the tank wall. How well a given choice of interaction zone ranges explains the data was estimated by the mean angle difference between its predicted and the actual direction after the burst. To find the parameter-choice for *r_r_*, *r_o_* and *r_a_* which minimizes this angle difference we, first, ran two different optimizer (dual-annealing: python/scipy implementation of (82), differential-evolution: python/scipy implementation of (83)) which gave for each selection line similar parameters. For all selection lines the width of the orientation zone was below 2 millimeters which suggests that the best solution favors no alignment at all. To verify this we did two-dimensional parameter scans around the optimal parameter setting (see Fig.S10). For the scan in which the repulsion *r_r_* and orientation range *r_o_* are varied, the angular difference is lowest if *r_o_* = *r_r_*, i.e. for a two-zone model without orientation zone(Fig.S10B, E, H). Consequently we set the *r_o_* = *r_r_* and did a parameter-scan (Fig.S11) around the optima of the selection lines. From this scan we extracted the ranges reported in Tab.1. Interestingly, the ranges of the large-harvested line are smaller than the ones of the random-harvested line, which again are smaller than the ones of the small-harvested line. Therefore, the ranges can be interpreted to represent body-length differences between the selection lines.

#### II. B. Parameter setting by fitting emergent variables

To fit the model to the data we applied the CMA-ES (Covariance Matrix Adaptation Evolution Strategy) (52, 53). This optimizer is a good choice if the fitness landscape is multi-modal, the search space dimension is between 4 and 100 and no gradient is known. The optimizer minimizes the error function

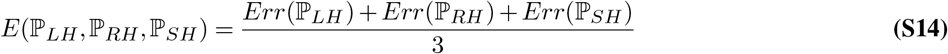

with 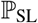 as the set of parameter of a selection line (SL) needed to run a simulation. Eq. S14 averages over the selection line errors. The error of a specific selection lines compares the measured nearest neighbour distance NND and average individual speed *v* to the experimental data by computing the sum of the squared differences standardized by their experimental standard deviation:

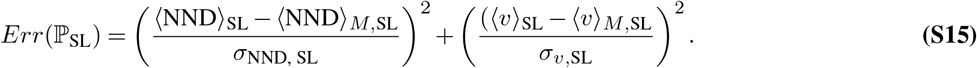

Here the index *M* in 〈x〉_*M*,SL_ marks that the measure of *x* is based on model simulations. 〈x〉_SL_ and *σ*_*x*,SL_ are the experimental mean and standard deviation of the measure *x*. SL specifies the selection line. The experimental means of NND and *v* used for the error function are listed in Tab. S2.

At each generation the parameter sets are updated by the CMA-ES method. Note that the parameter sets differ in burst-duration, -rate, repulsion-, attraction-range and probability to respond to social cues in between the selection lines. Only the burst force *F* and the probability to respond to social cues need to be set. Consequently, the search-space is 6-dimensional (*P_soc,LH_*, *P_soc,RH_*, *P_soc,SH_*, *F_LH_*, *F_RH_*, *F_SH_*) as highlighted in Tab. 1.

We limited the search-space of the algorithm for the three different parameters by setting boundaries which are listed in Tab. S2. For the probability to follow social cues we ensured a minimum attention to social and environmental cues by setting the boundaries 0.05 above and below the theoretical possible boundaries of zero and 1. We expect the burst force to be larger than the experimentally estimated measure. Therefore, it’s boundaries are half and twice the mean burst force of the random harvested line, estimated in section II. A.

To ensure that the resulting minimum is not a local minimum, we repeated the optimization from different initial parameter settings. The initial parameter were selected from a two-dimensional grid with 2 grid-points and therefore 2^2^ =4 different initial parameter settings. Note that the actual search-space is 6-dimensional and by setting *P_soc_* and *F* for the different selection line initially equal we reduce the number of initial settings from 2^6^ = 64 to 4.

An example optimization run over 400 generations is shown in Fig. S8. The optimization outcomes of the 4 different initialization are shown sorted by their final error according to Eq. S14 in Fig. S9.

### III Mean distance of shoaling fish to fishing agents

Here we computed the mean distance the shoaling fish kept from fishing agents. This measure is an alternative to the mortality rate used in the main text, to quantify how well the shoaling fish avoid capture:

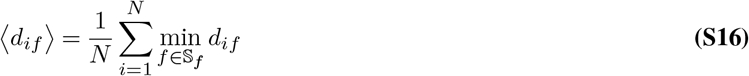

with 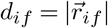 as the distance between shoaling fish *i* and a fishing agent *f*. The minimum function yields the distance to the closest fishing agent. In Fig. S3I, J, K, L show the relative and M, N, O, P the absolute distance to the predator. The distance measured substantiated the already reported results for the mortality rate. Note that in the natural-predator scenario (I, M) the differences between selection lines is qualitatively equivalent to the other scenarios in contrast to the scenario comparison with the mortality rate(A, E). This is due to (i) the confusion effect, the main difference between the natural predator and the other fishing agents, which reduces the mortality rate for more cohesive shoals (RH and SH line) but does not affect the distance and (ii) the fast individual speed of the RH line, which enables the currently pursued fish to better escape but has no effect on how the other shoaling agents detect and avoid the predator. Note that the natural predator moves at constant speed and is therefore only on average faster than the shoaling fish.

### IV Movies

**Movie-S1.avi.** Eight individuals are simulated in a circular area with fitted social interaction. The parameters were the ones estimated for the random harvested line listed in Tab. 1.

**Movie-S2.avi.** This movie shows simulations in the circular area with parameters adjusted such that only the wall avoidance mechanism changes the path of the shoaling fish. The parameters are the ones estimated for the random harvested line listed in Tab. 1 apart for the repulsion, alignment and attraction range and probability to follow social cues *P_soc_*. Those parameters were set such that the agents always responded to social information *P_soc_* = 1, but never had any neighbors due to short ranges *r_r_* = *r_o_* = *r_a_* = 0.001*cm*. Thus, the agents effectively swim always straight until they come in conflict with the wall. Then the wall avoidance mechanism started.

**Movie-S3.avi.** Thirty agents simulated in a rectangle of length *L* = 100*cm* with periodic boundary conditions in the predator scenario. The parameters were the ones estimated for the random harvested line listed in Tab. 1 and additional fishing/predator scenario parameters listed in Tab. S3. Note that the predator, in red, moves with a speed of *v_p_* = 20*cm/s* and therefore faster as the average velocity of shoaling fish. However, it moves directly to the closest prey and therefore covers the prey until it get killed. Due to the confusion effect the killing probability was lowered and it happens that the predator stays for a notable time above the targeted fish.

**Movie-S4.avi.** Thirty agents simulated in a rectangle of length *L* = 100*cm* with periodic boundary conditions in the angler scenario. The parameters were the ones estimated for the random harvested line listed in Tab. 1 and additional fishing/predator scenario parameters listed in Tab. S3.

**Movie-S5.avi.** Thirty agents simulated in a rectangle of length *L* = 100*cm* with periodic boundary conditions in the trawling scenario. The parameters were the ones estimated for the random harvested line listed in Tab. 1 and additional fishing/predator scenario parameters listed in Tab. S3.

**Fig.S1.**
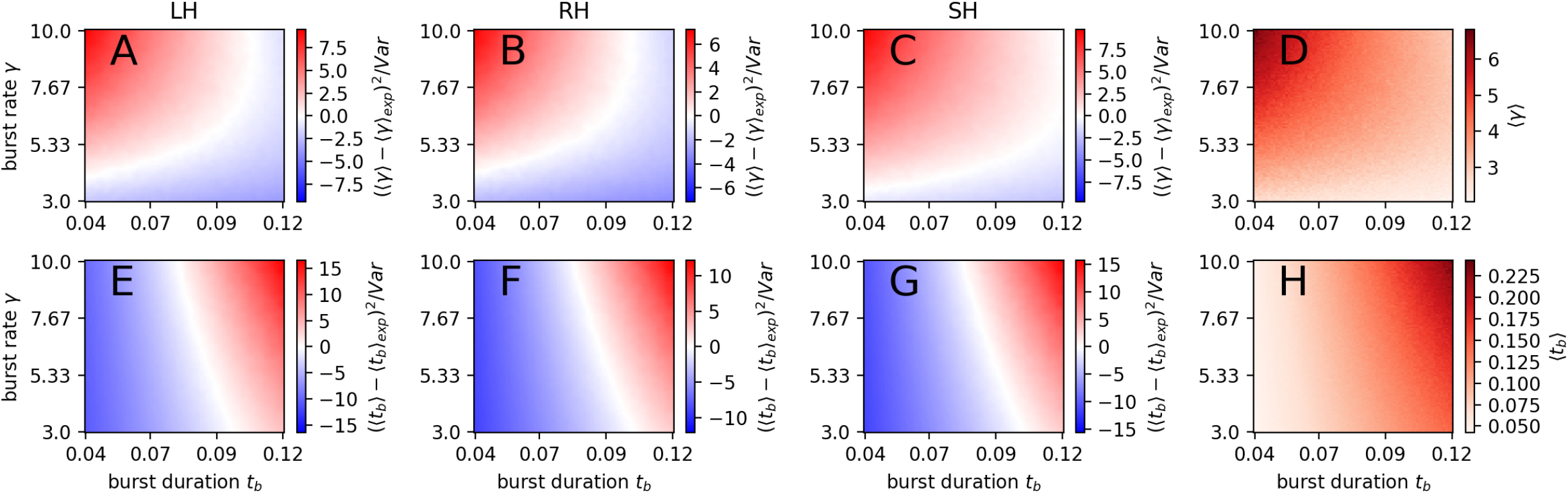
Model parameter estimation of burst duration *t_b_* and burst rate *γ* (parameters of an acceleration-time-series generating model). The acceleration-based-average is reported for burst rate 〈*γ*〉 (**A-D**) and burst duration (**E-H**) 〈*t_b_*〉. The normalized squared difference between the averaged values of the simulation 〈*x*〉 and the experiment 〈*x*〉_*exp*_ are shown for large-harvested line (LH; **A, E**), random-harvested line (RH; **B, F**) and small-harvested line (SH;**C, G**). The normalization was done by dividing the squared differences by the experimental variance of the burst rate (**A-C**) or of the burst duration (**E-G**). The average values of the simulation are shown in (**D, H**).

**Fig. S2.**
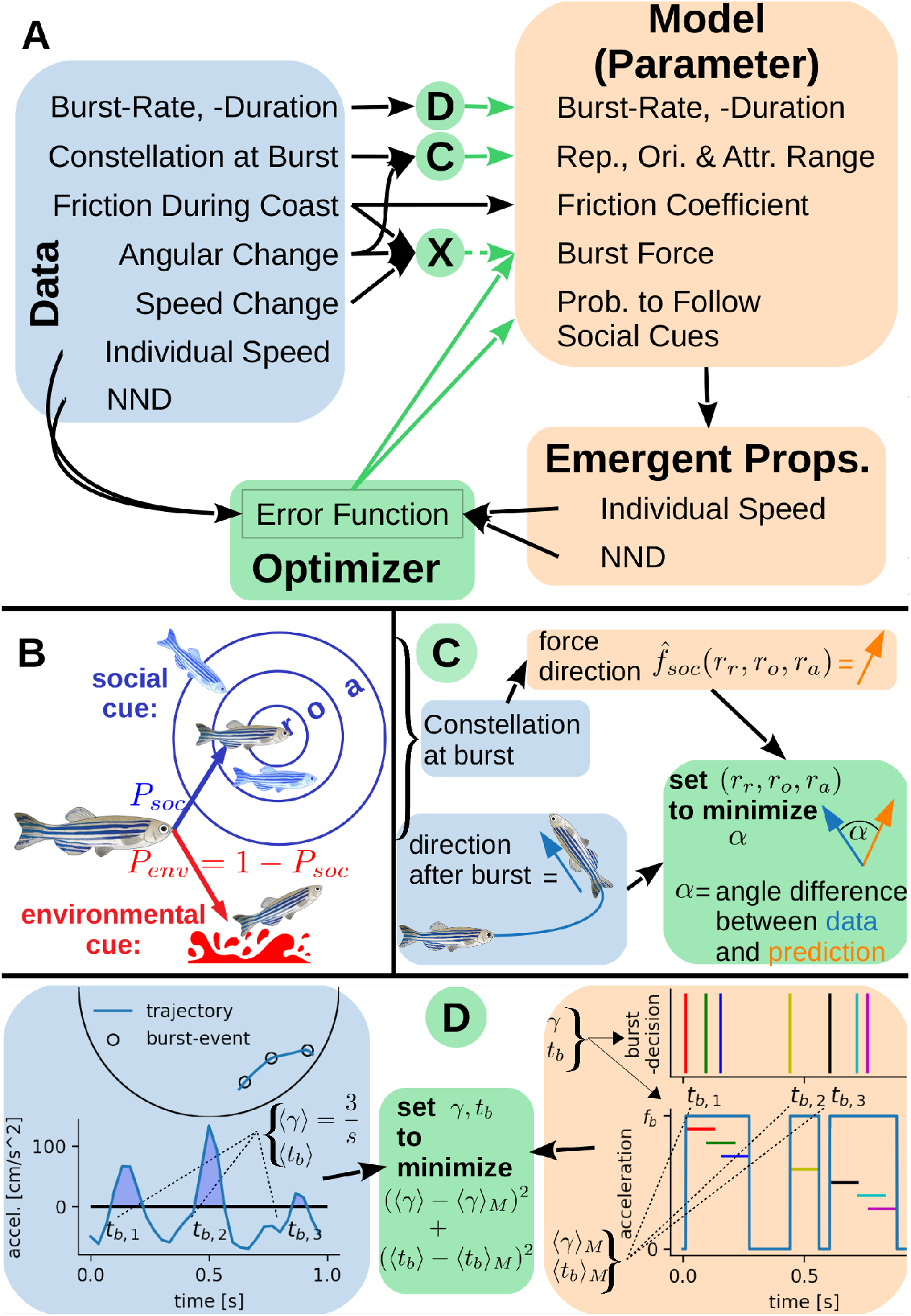
Summary of model parameter setting from data. **A**: relations between trajectory data based measurements and model parameters. All light green boxes/circles represent optimizer which minimize an error-function by adjusting the model parameters. The model parameter setting is indicated by green arrows. The central green circle marked with an “X” represents the optimizer which deduces the burst force parameter. However, since our estimation of the burst force *F* is likely to be too low (as explained in SI Sect. II. A) and therefore is estimated by the optimizer which compares NND “and speed, we omit an illustrative explanation of its dependence. **B**: model scheme of burst decisions. An individual can follow with probability *P_soc_* social cues (i.e. behaves according to a three-zone model: r=repulsion-, o=orientation-, a=attraction-zone) or with probability *P_env_* = 1 – *P_soc_* environmental cues. **C**: sketch of how the range of repulsion *r_r_*, orientation *r_o_*, and attraction *r_a_* are set by minimizing the angle difference *α* between the model-predicted force direction 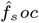 and the actual heading direction after a burst. **D**: setting of the burst rate *γ* and duration *t_b_*. The difference between the averages based on data (〈*γ*〉, 〈*t_b_*〉 blue box) and on model-simulations (〈*γ*〉_*M*_, 〈*t_b_*〉_*M*_ orange box) is minimized.

**Fig. S3.**
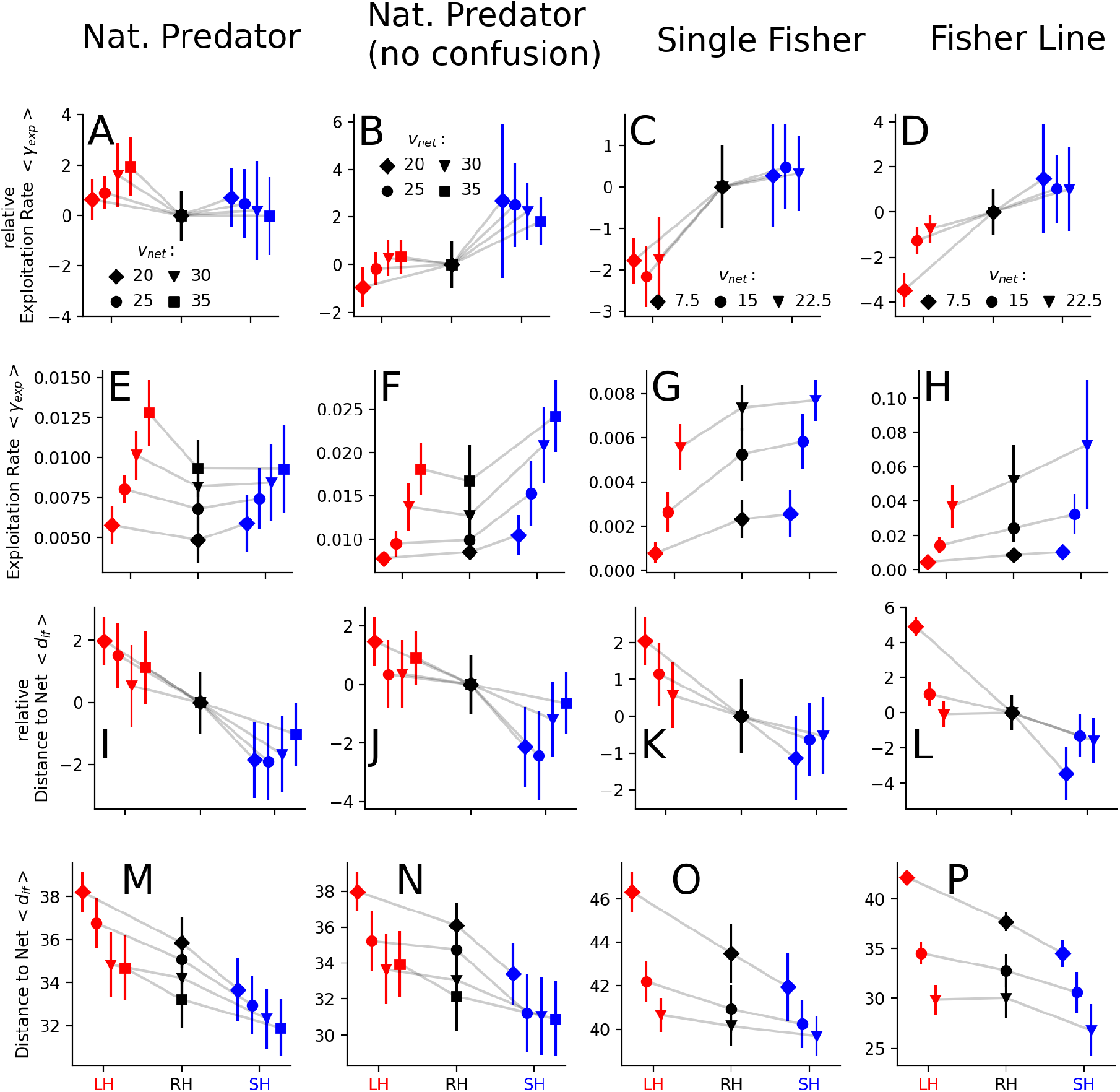
The relative (**A-D**) and normal (**E-H**) mean mortality rate *γ_exp_* and the relative (**I-L**) and normal (**M-P**) mean distance to the closest fishing agent *d_if_* are shown for different scenarios. The relative measures were computed by reducing the selection line means by the mean and dividing them by the standard deviation of the random-harvested line. In the “natural predator” scenario a predator pursues the closest shoaling fish with different speeds *v_net_* = [20, 25, 30, 35] *cm/s* but its capture success is reduced the more prey it senses, i.e. by the confusion effect (1st column, **A, E, I, M**). The “natural predator without confusion” scenario is the same but without the confusion effect (2nd column, **B, F, J, N**). In the “single fisher” scenario an uninformed single fisher moves randomly and captures shoaling agents at encounter (3rd column, **C, G, K, O**). In the “multiple fishing agent” scenario fishing agents move *N_f_* = 50 fishing agent on a rigid line towards the center of mass of the fish shoal (4th column, **D, H, L, P**). The speed of the fishing agents in the “single fisher” and “mutliple fishing agent” scenarios was *v_net_* = [7.5, 15, 22.5] *cm/s* indicated by rhombus, circle, triangle, respectively. In all simulations *N* = 30 shoaling fish moved in a box of size *L* = 100 *cm* with periodic boundary conditions. Their mean individual speed was about 〈*v*〉 ≈ 15 *cm/s*. The symbols and error bars represent the mean and standard deviation computed from samples of 400 simulations.

**Table S2.** The upper part of the table lists the upper and lower boundaries in which the optimizer searched for the minimum of the error function Eq. S14. The lower and upper boundary of the burst force *F* are given by half and twice the mean of the experimental estimated burst force 〈*f_b_*〉_*RH*_ of the RH line. The lower part of the table lists the measures defining the error function Eq. S14 which is minimized by the optimizer.

| optimization parameter | symbol | optimization boundaries | unit |
| --- | --- | --- | --- |
| burst force | $F$ | $[67.6, 270.4]$<br>($= [\langle f_b \rangle_{RH}/2, 2\langle f_b \rangle_{RH}]$) | $cm/s^2$ |
| prob. of social burst | $P_{soc}$ | $[0.05, 0.95]$ | 1 |
| error function defining measures | symbol | experimental mean and STD | unit |
| individual speed | $v$ | LH: $15 \pm 2.8$<br>RH: $17 \pm 2.5$<br>SH: $13.2 \pm 2.7$ | $cm/s$ |
| nearest neighbor dist. | $NND$ | LH: $6.4 \pm 1$<br>RH: $5.8 \pm 1.1$<br>SH: $5.6 \pm 0.9$ | $cm$ |

**Table S3.** Parameters used to simulate interaction of fish with predator/fisher in different scenarios. The column “scenario relevance” specifies for which scenario the parameter is needed. ^*a*^: Note that the flee range influences the probability of a prey detecting a predator/fisher (Eq. S6) which is 0.5 at a distance of 21*cm* given *r_f_* = 7*cm*. Therefore, the range of the prey to detect a predator is comparable to the predators sensing range.

| scenario related parameter | symbol | value | scenario relevance | unit |
| --- | --- | --- | --- | --- |
| flee range<sup>a</sup> | $r_f$ | 7 | all | $cm$ |
| fishers/predators capture range | $r_{capture}$ | 5 | all | $cm$ |
| predators sensing range | $r_{sense}$ | $4r_{capture} = 20$ | natural pred. | $cm$ |
| confusion number | $N_{conf}$ | 4 | natural pred. | 1 |
| attack rate | $\gamma_a$ | 1 | natural pred. | $1/s$ |
| angular noise strength | $\sigma_\varphi$ | 0.2 | single fisher | $1/s^{1/2}$ |

**Fig. S4.**
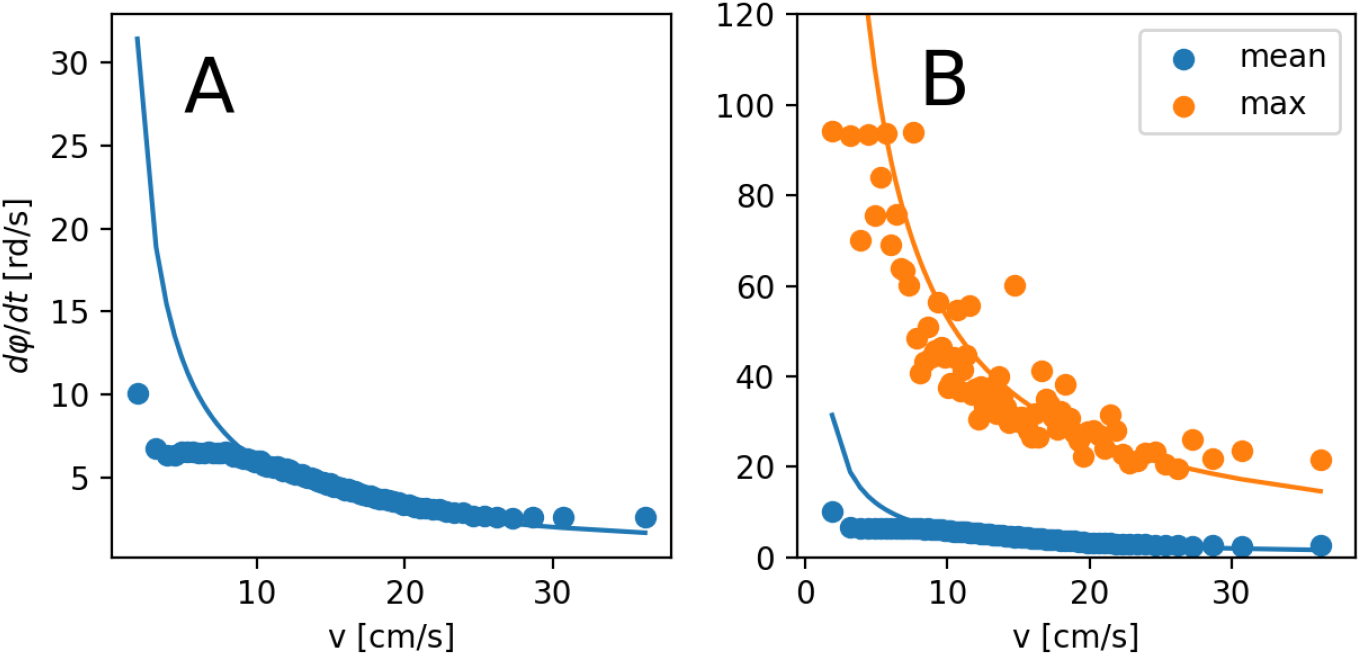
Angular change *dφ/dt* depending on the velocity *v*. The blue dots (**A, B**) represent the mean values while orange dots (**B**) the maximum observed value for the binned data. The lines are not fitted to the data but serve as an comparison to the inverse proportionality. The blue line is *f* (*v*) = 6/*v* and the orange line is *f* (*v*) = 530*/v*. The resemblance of the data to the lines supports the model assumption in Eq. (S2c).

**Fig. S5.**
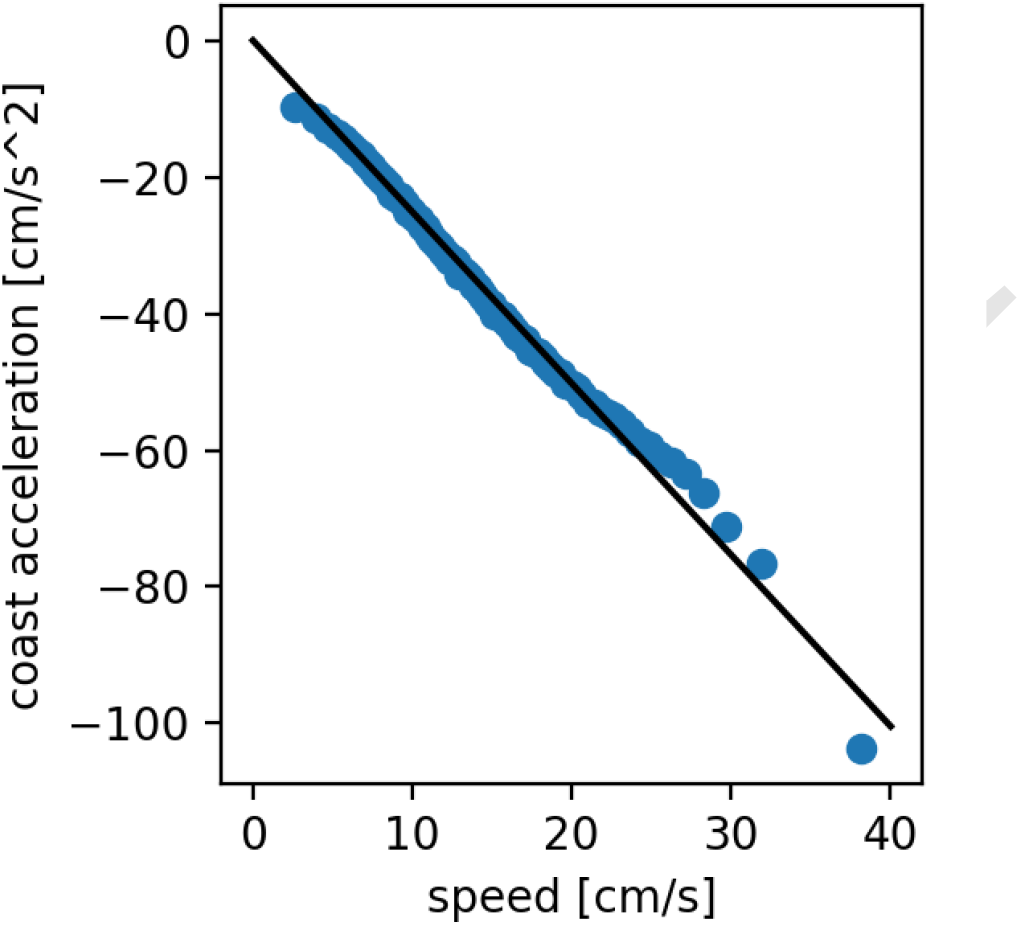
Estimation of friction coefficient from experimental data. Dots represent mean of bins with varying width such that each bin contains an equal amount of data. Black line shows linear fit with the interception at zero and the negative friction coefficient as slope – *β* = −2.51.

**Fig. S6.**
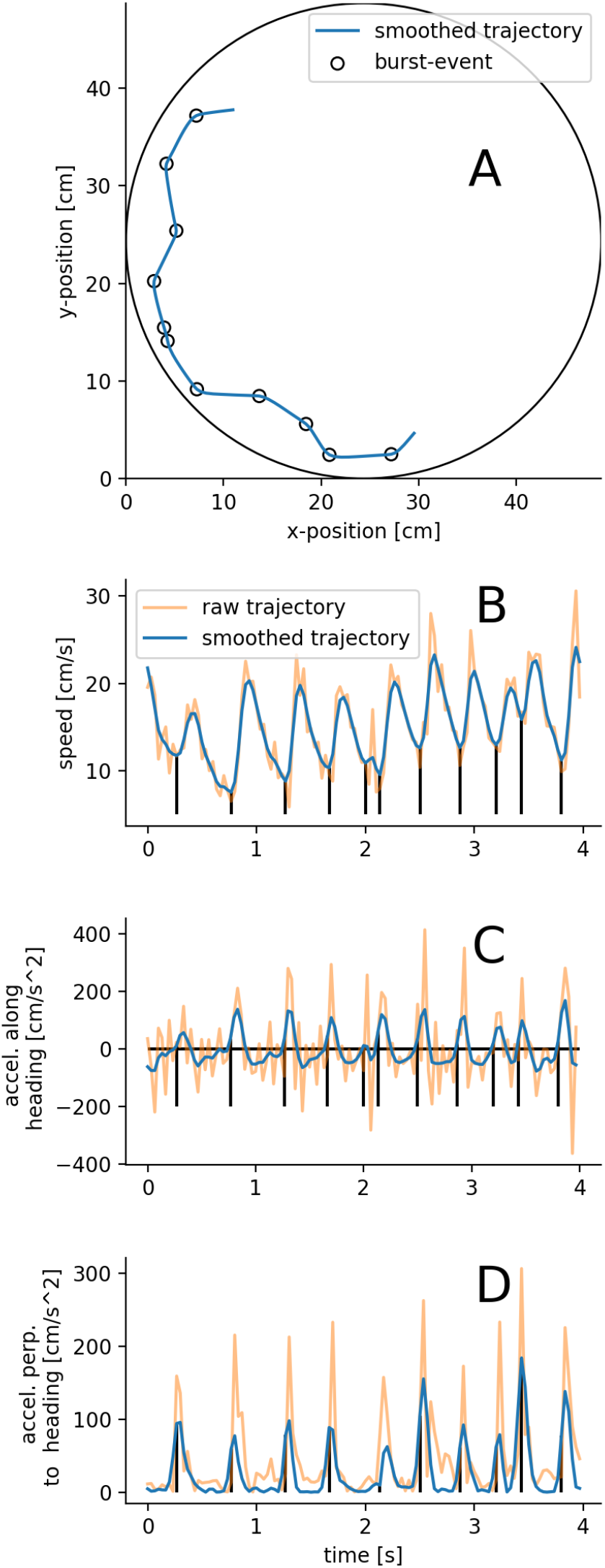
Example smoothed trajectory of a swimming zebrafish generated from ID-Tracker data (**A**). The large circle represents tank-boundary. For this trajectory the speed (**B**), acceleration along (**C**) and perpendicular (**D**) to heading direction are shown computed from the raw (orange) and smoothed (blue) trajectory. The black circles (**A**) and vertical black lines (**B, C, D**) indicate burst events which are are the onsets of a positive acceleration period of the smoothed data (**C**). For the smoothening a moving window average with gaussian kernel was used. The kernel width *σ_smoo_* = 1.13*fms* was estimated by comparing manual counts of burst events with automatically detected (see Fig. S7).

**Fig. S7.**
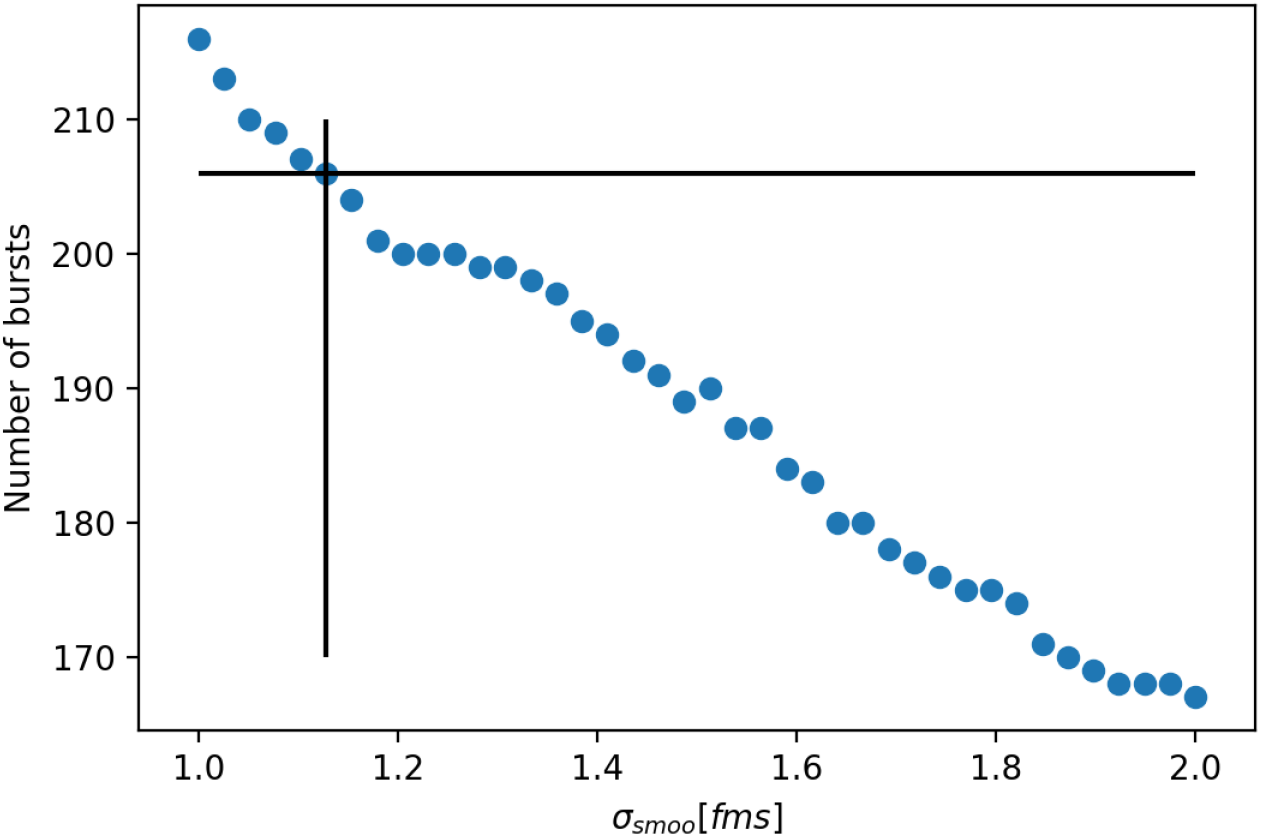
Number of burst events of a randomly selected trajectory spanning 2000 frames. The horizontal black line marks the number of manually counted burst events. Blue dots mark automatic counts of burst events on a smoothed trajectory using a moving window with a Gaussian kernel of varying standard deviation *σ_smoo_*. The vertical black line indicates the standard deviation *σ_smoo_* = 1.13 *fms* where the number of automatically detected bursts equals the manually counted one in an interval of 2000 frames.

**Fig. S8.**
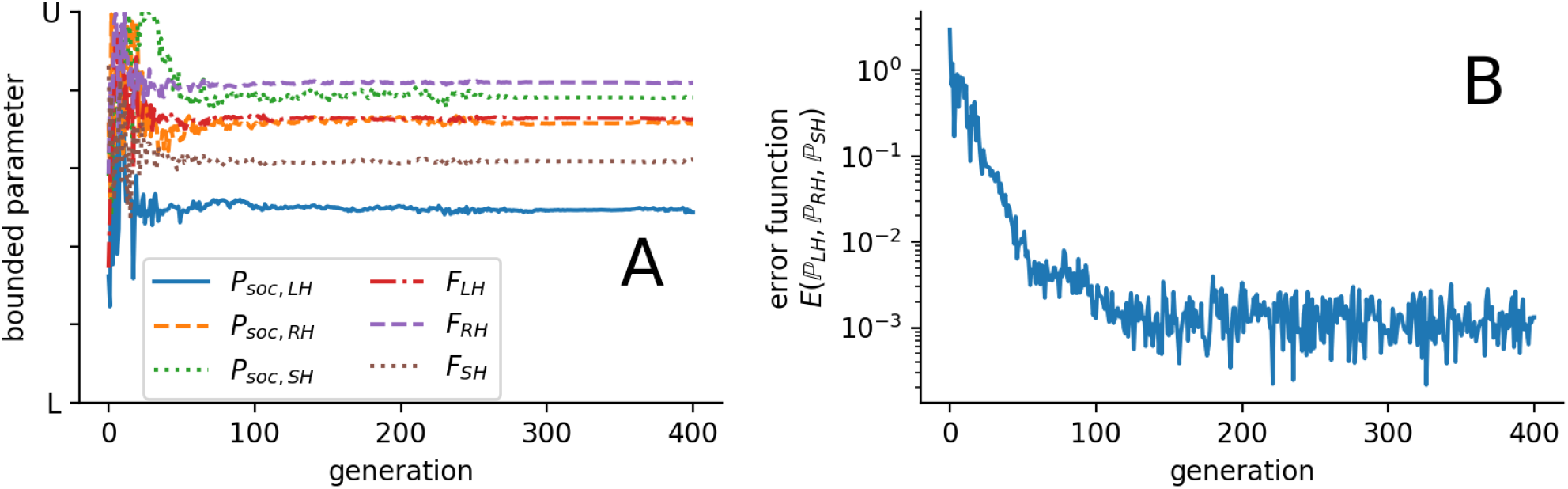
Example of optimization run with the parameters which result in the lowest error in the current generation (**A**) and the corresponding error (**B**). The parameters are normalized according to their respective lower and upper boundary (see Tab. S2) which are marked on the y-axis with L and U (**A**). Different colors and linestyles mark the probability to respond to social cues *Psoc* of the large (solid blue), random (dashed orange) and small (dotted green) harvested line and the burst forces *F* of the large (dash dotted red), random (dashed violet) and small (dotted brown) harvested line.

**Fig. S9.**
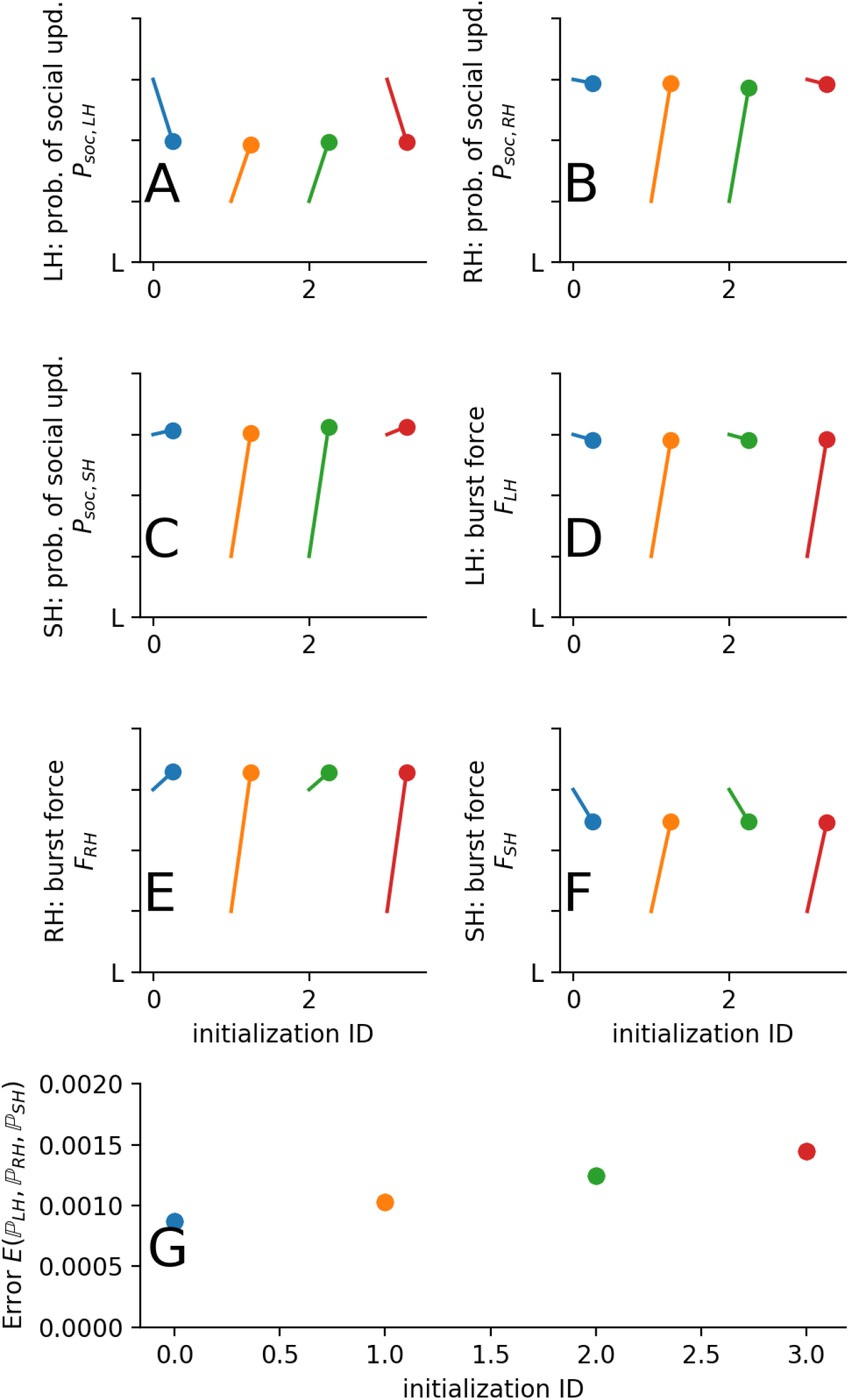
Initial and final parameters (**A-F**) together with final error (**G**) according to Eq. S14 for the 4 different initialized optimization runs. The initialization are sorted such that the one with the best outcome, i.e. with the smallest error, is first. Colors mark specific initialization ID and are consistent across subfigures. The circles (**A-E**) mark the final parameter while the other end of the line marks the initial setting.

**Fig. S10.**
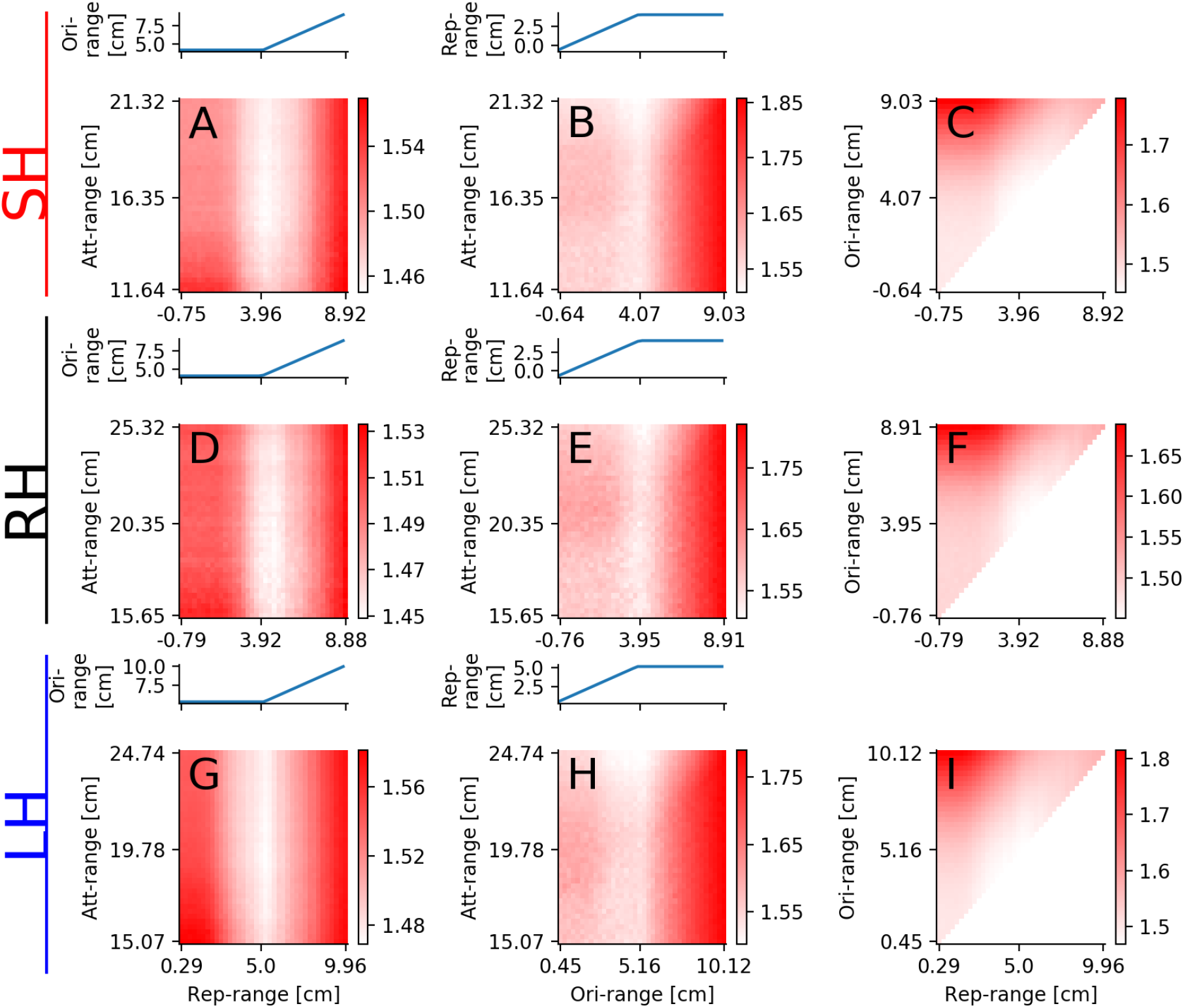
**Angle difference** between predicted and actual direction afterburst is color-coded. The difference is analyzed around the optimal values of the repulsion, orientation and attraction ranges. The line-plots above the color-plots show the remaining range which was kept constant if possible but needed to be in- or decreased to ensure the inequality *r_r_* < *r_o_* < *r_a_*. In the first (**A, B, C**), second (**D, E, F**) and third (**G, H, I**) row is data used from burst-constellations of the small-, random- and large-harvested selection line, respectively. We varied the repulsion and attraction range in the first column (**A, D, G**), the orientation and attraction range in the second (**B, E, H**) and the repulsion and orientation range in the third column (**C, F, I**).

**Fig. S11.**
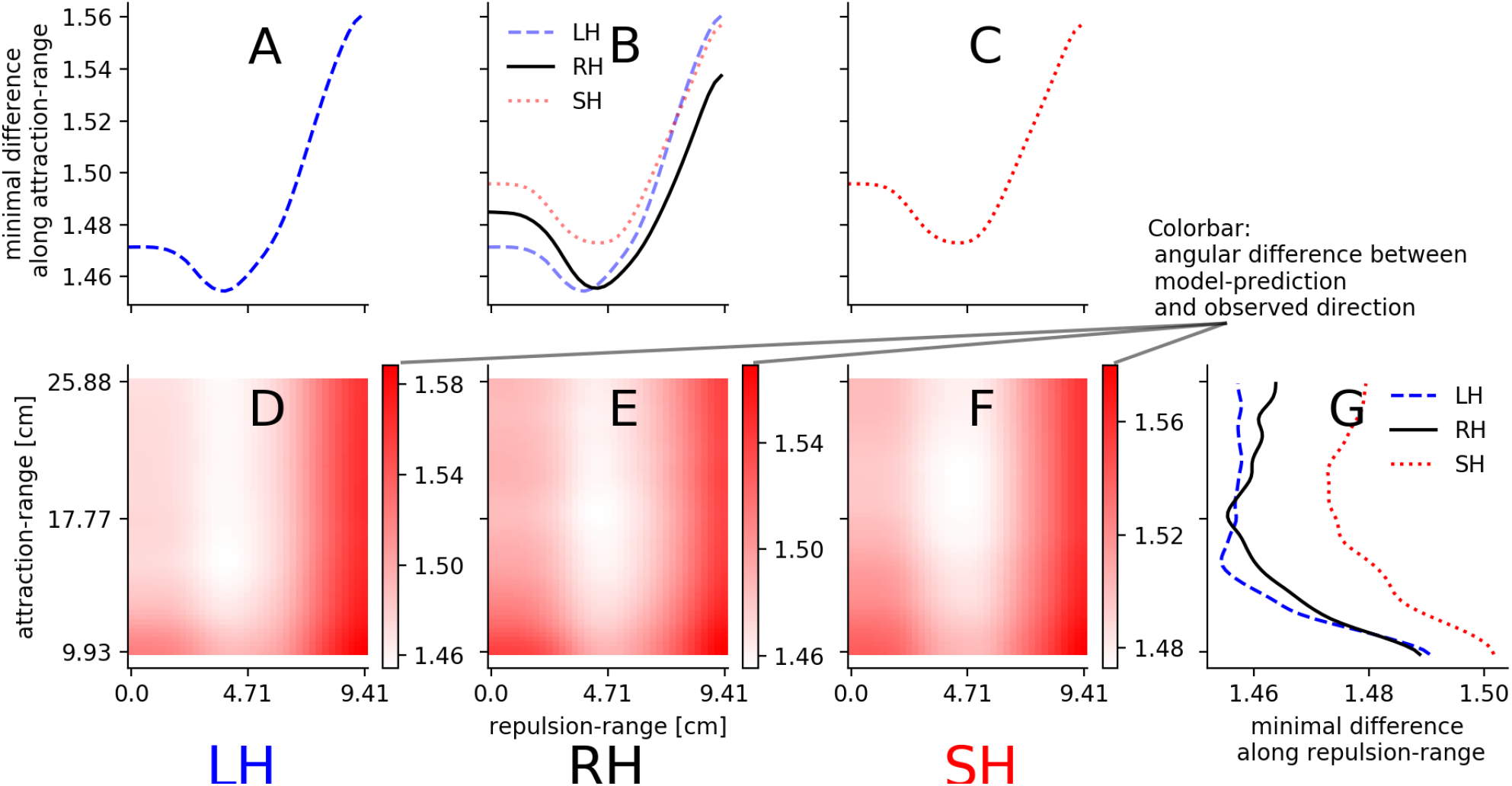
**Angle difference** between predicted and actual direction after burst is color-coded (**D-F**) or its minimal value along one axis direction is shown (**A-C, G**). Here, in contrast to the scans in Fig.S10, no orientation-zone exists, i.e. *r_o_* = *r_r_*. **A-C**: minimal angle difference along the attraction-range is shown for burst-constellation data of the large- (**A**), small-harvest (**C**) and for all lines (**B**). **D-F**: angular difference for different parameters of repulsion and attraction range for the large-(**D**), random-(**E**) and small-harvested line(**F**). **G**: minimal difference for a specific attraction-range for all selection lines.

**Fig. S12.**
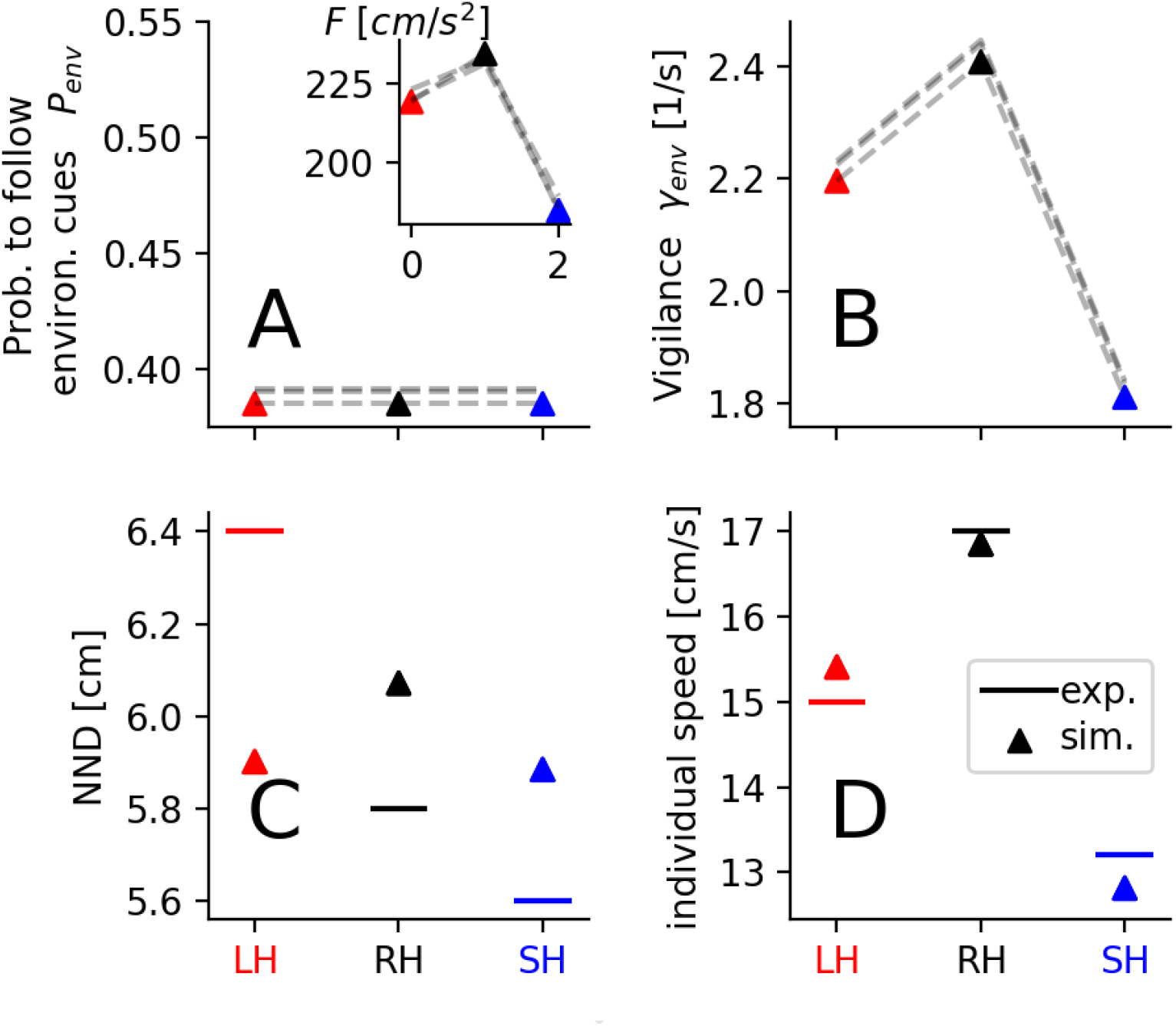
Fitted model representation of the different selection lines. Note that here, in **contrast to Fig. 4 in the main text, we enforced the probability to follow environmental cues to be the same for all selection lines**. In order to resemble the experimental observations the probability to follow environmental cues (**A**) and the burst force (inset **A**) were set for each selection line. The vigilance *γ_env_* is the product of *P_env_* and the burst rate *γ* (**B**). The average nearest neighbor distance (NND, **C**) and the average individual speed (**D**) are emergent properties of the model and were used to quantify how well the parameters (*P_env_, γ*) reproduce the experimental observations. In all panels: the triangles represent to the model parameters or the simulation outcomes of the parameter set with the best match to the experiment. Each dashed line (**A, B**) represents a parameter set of a different initialization, and therefore illustrates the robustness of the best matching parameter set (triangles). The horizontal solid lines (**C, D**) mark the experimental values. Note that in contrast to Fig. 3 units are in *cm* because the modelled agents have no body length. The selection lines are indicated by different colors (red=LH: large-harvested; black=RH: random-harvested; blue=SH: small-harvested).

## Bibliography

1. Susan M Sogard. Size-selective mortality in the juvenile stage of teleost fishes: a review. Bulletin of marine science, 60(3):1129–1157, 1997. ISSN 0007-4977.

2. Kai Lorenzen. Allometry of natural mortality as a basis for assessing optimal release size in fish-stocking programmes. Canadian Journal of Fisheries and Aquatic Sciences, 57(12):2374–2381, 2000. ISSN 0706-652X. doi: 10.1139/f00-215.

3. Mikko Heino, Beatriz Díaz Pauli, and Ulf Dieckmann. Fisheries-induced evolution. Annual Review of Ecology, Evolution and Systematics, 46(1):461–480, 2015. ISSN 1543-592X. doi: 10.1146/annurev-ecolsys-112414-054339.

4. David O. Conover and Stephan B. Munch. Sustaining fisheries yields over evolutionary time scales. Science, 297(5578):94–96, 2002. doi: 10.1126/science.1074085.

5. Silva Uusi-Heikkilä, Andrew R. Whiteley, Anna Kuparinen, Shuichi Matsumura, Paul A. Venturelli, Christian Wolter, Jon Slate, Craig R. Primmer, Thomas Meinelt, Shaun S. Killen, David Bierbach, Giovanni Polverino, Arne Ludwig, and Robert Arlinghaus. The evolutionary legacy of size-selective harvesting extends from genes to populations. Evolutionary Applications, 8(6):597–620, 2015. ISSN 1752-4571. doi: 10.1111/eva.12268.

6. Ken H. Andersen, Lise Marty, and Robert Arlinghaus. Evolution of boldness and life history in response to selective harvesting. Canadian Journal of Fisheries and Aquatic Sciences, 75(2):271–281, 2018. ISSN 0706-652X 1205-7533. doi: 10.1139/cjfas-2016-0350.

7. V. Sbragaglia, J. F. Lopez-Olmeda, E. Frigato, C. Bertolucci, and R. Arlinghaus. Size-selective mortality induces evolutionary changes in group risk-taking behaviour and the circadian system in a fish. J Anim Ecol, 90(2):387–403, 2021. ISSN 1365-2656 (Electronic) 0021-8790 (Linking). doi: 10.1111/1365-2656.13372.

8. Darren P Croft, Jens Krause, Iain D Couzin, and Tony J Pitcher. When fish shoals meet: outcomes for evolution and fisheries. Fish and Fisheries, 4(2):138–146, 2003. ISSN 1467-2960. doi: 10.1046/j.1467-2979.2003.00113.x.

9. Ana Sofia Guerra, Albert B Kao, Douglas J McCauley, and Andrew M Berdahl. Fisheries-induced selection against schooling behaviour in marine fishes. Proc. R. Soc. B Biol. Sci., 287(1935):20201752, sep 2020. ISSN 0962-8452. doi: 10.1098/rspb.2020.1752.

10. Robert Arlinghaus, Kate L. Laskowski, Josep Alós, Thomas Klefoth, Christopher T. Monk, Shinnosuke Nakayama, and Arne Schröder. Passive gear-induced timidity syndrome in wild fish populations and its potential ecological and managerial implications. Fish and Fisheries, 18(2):360–373, 2017. ISSN 1467-2979. doi: 10.1111/faf.12176.

11. W. D. Hamilton. Geometry for the selfish herd. Journal of Theoretical Biology, 31(2):295–311, 1971. ISSN 0022-5193. doi: https://doi.org/10.1016/0022-5193(71)90189-5.

12. Jens Krause and Graeme D Ruxton. Living in groups. Oxford University Press, 2002. ISBN 0198508182.

13. Ashley Ward and Mike Webster. Sociality: the behaviour of group-living animals. 2016.

14. Manfred Milinski. Experiments on the selection by predators against spatial oddity of their prey. Zeitschrift für Tierpsychologie, 43(3):311–325, 1977. ISSN 0044-3573. doi: 10.1111/j.1439-0310.1977.tb00078.x.

15. Tony J. Pitcher, Ole Arve Misund, Anders Fernö, Bjørn Totland, and Vebjørn Melle. Adaptive behaviour of herring schools in the norwegian sea as revealed by high-resolution sonar. ICES Journal of Marine Science, 53(2):449–452, 1996. ISSN 1054-3139. doi: 10.1006/jmsc.1996.0063.

16. Timothy M. Schaerf, Peter W. Dillingham, and Ashley J. W. Ward. The effects of external cues on individual and collective behavior of shoaling fish. Science Advances, 3(6):e1603201, 2017. doi: 10.1126/sciadv.1603201.

17. S W Milne, B J Shuter, and W G Sprules. The schooling and foraging ecology of lake herring (coregonus artedi) in lake opeongo, ontario, canada. Canadian Journal of Fisheries and Aquatic Sciences, 62(6):1210–1218, 2005. doi: 10.1139/f05-030.

18. Jens Krause and Jean-Guy J. Godin. Influence of prey foraging posture on flight behavior and predation risk: predators take advantage of unwary prey. Behavioral Ecology, 7(3):264–271, 1996. ISSN 1045-2249. doi: 10.1093/beheco/7.3.264.

19. Hannah E. A. MacGregor, James E. Herbert-Read, and Christos C. Ioannou. Information can explain the dynamics of group order in animal collective behaviour. Nature Communications, 11(1):2737, dec 2020. ISSN 2041-1723. doi: 10.1038/s41467-020-16578-x.

20. P. Domenici and R. S. Batty. Escape behaviour of solitary herring (Clupea harengus) and comparisons with schooling individuals. Marine Biology, 128(1):29–38, apr 1997. ISSN 0025-3162. doi: 10.1007/s002270050065.

21. Guy Beauchamp. Group-size effects on vigilance: A search for mechanisms. Behavioural Processes, 63(3):111–121, 2003. ISSN 03766357. doi: 10.1016/S0376-6357(03)00002-0.

22. Guy Beauchamp. What is the magnitude of the group-size effect on vigilance? Behavioral Ecology, 19(6):1361–1368, 2008. ISSN 10452249. doi: 10.1093/beheco/arn096.

23. Ashley J.W. Ward, James E. Herbert-Read, David J.T. Sumpter, and Jens Krause. Fast and accurate decisions through collective vigilance in fish shoals. Proceedings of the National Academy of Sciences of the United States of America, 108(6):2312–2315, 2011. ISSN 10916490. doi: 10.1073/pnas.1007102108.

24. Sybille Hess, Stefan Fischer, and Barbara Taborsky. Territorial aggression reduces vigilance but increases aggression towards predators in a cooperatively breeding fish. Animal Behaviour, 113:229–235, 2016. ISSN 00033472. doi: 10.1016/j.anbehav.2016.01.008.

25. Julia K. Parrish. Using behavior and ecology to exploit schooling fishes. Environmental Biology of Fishes, 55(1):157–181, 1999. ISSN 1573-5133. doi: 10.1023/A:1007472602017.

26. Alexander Kotrschal, Alexander Szorkovszky, James Herbert-Read, Natasha I. Bloch, Maksym Romenskyy, Séverine Denise Buechel, Ada Fontrodona Eslava, Laura Sánchez Alòs, Hongli Zeng, Audrey Le Foll, Ganaёl Braux, Kristiaan Pelckmans, Judith E. Mank, David Sumpter, and Niclas Kolm. Rapid evolution of coordinated and collective movement in response to artificial selection. bioRxiv, page 2020.01.30.926311, 2020. doi: 10.1101/2020.01.30.926311.

27. Daniel S. Calovi, Alexandra Litchinko, Valentin Lecheval, Ugo Lopez, Alfonso Pérez Escudero, Hugues Chaté, Clément Sire, and Guy Theraulaz. Disentangling and modeling interactions in fish with burst-and-coast swimming reveal distinct alignment and attraction behaviors. PLOS Computational Biology, 14(1):e1005933, jan 2018. ISSN 1553-7358. doi: 10.1371/journal.pcbi.1005933.

28. Iain D. Couzin and Jens Krause. Self-Organization and Collective Behavior in Vertebrates, volume 32, pages 1–75. Academic Press, 2003. ISBN 0065-3454. doi: https://doi.org/10.1016/S0065-3454(03)01001-5.

29. James E Herbert-Read, Andrea Perna, Richard P Mann, Timothy M Schaerf, David JT Sumpter, and Ashley JW Ward. Inferring the rules of interaction of shoaling fish. Proceedings of the National Academy of Sciences, USA, 108(46):18726–18731, 2011. ISSN 0027-8424.

30. Iain D Couzin, Jens Krause, Richard James, Graeme D Ruxton, and Nigel R Franks. Collective memory and spatial sorting in animal groups. Journal Theoretical Biology, 218(1):1–11, 2002. ISSN 00225193. doi: 10.1006/yjtbi.3065.

31. Iain D Couzin, Christos C Ioannou, Güven Demirel, Thilo Gross, Colin J Torney, Andrew Hartnett, Larissa Conradt, Simon A Levin, and Naomi E Leonard. Uninformed Individuals Promote Democratic Consensus in Animal Groups. Science, 334(6062):1578–1580, dec 2011. ISSN 0036-8075. doi: 10.1126/science.1210280.

32. J. Buhl, D. J. T. Sumpter, I. D. Couzin, J. J. Hale, E. Despland, E. R. Miller, and S. J. Simpson. From Disorder to Order in Marching Locusts. Science, 312(5778):1402–1406, jun 2006. ISSN 0036-8075. doi: 10.1126/science.1125142.

33. Jolle W. Jolles, Andrew J. King, and Shaun S. Killen. The role of individual heterogeneity in collective animal behaviour. Trends in Ecology & Evolution, 2019. ISSN 0169-5347. doi: https://doi.org/10.1016/j.tree.2019.11.001.

34. Noam Miller and Robert Gerlai. From schooling to shoaling: patterns of collective motion in zebrafish (danio rerio). PLoS One, 7(11):e48865, 2012. ISSN 1932-6203.

35. R. Spence, G. Gerlach, C. Lawrence, and C. Smith. The behaviour and ecology of the zebrafish, danio rerio. Biol Rev Camb Philos Soc, 83(1):13–34, 2008. ISSN 1464-7931 (Print) 0006-3231. doi: 10.1111/j.1469-185X.2007.00030.x.

36. Piyumika S Suriyampola, Delia S Shelton, Rohitashva Shukla, Tamal Roy, Anuradha Bhat, and Emilia P Martins. Zebrafish social behavior in the wild. Zebrafish, 13(1):1–8, 2016. ISSN 1545-8547.

37. N. Speedie and R. Gerlai. Alarm substance induced behavioral responses in zebrafish (*Danio rerio*). Behav Brain Res, 188(1):168–77, 2008. ISSN 0166-4328 (Print) 0166-4328. doi: 10.1016/j.bbr.2007.10.031.

38. Roy Harpaz and Elad Schneidman. Social interactions drive efficient foraging and income equality in groups of fish. Elife, 9:e56196, 2020. ISSN 2050-084X.

39. Roy Harpaz, Gašper Tkačik, and Elad Schneidman. Discrete modes of social information processing predict individual behavior of fish in a group. Proceedings of the National Academy of Sciences, USA, 114(38):10149–10154, 2017. ISSN 1091-6490 0027-8424. doi: 10.1073/pnas.1703817114.

40. Silva Uusi-Heikkilä, Kai Lindström, Noora Parre, Robert Arlinghaus, Josep Alós, and Anna Kuparinen. Altered trait variability in response to size-selective mortality. Biology Letters, 12(9):20160584, 2016. doi: 10.1098/rsbl.2016.0584.

41. S. Uusi-Heikkilä, T. Savilammi, E. Leder, R. Arlinghaus, and C. R. Primmer. Rapid, broad-scale gene expression evolution in experimentally harvested fish populations. Molecular Ecology, 26(15):3954–3967, 2017. ISSN 1365-294X (Electronic) 0962-1083 (Linking). doi: 10.1111/mec.14179.

42. Valerio Sbragaglia, Josep Alós, Kim Fromm, Christopher T. Monk, Carlos Díaz-Gil, Silva Uusi-Heikkilä, Andrew E. Honsey, Alexander D.M. Wilson, and Robert Arlinghaus. Experimental size-selective harvesting affects behavioral types of a social fish. Transactions of the American Fisheries Society, 148(3):552–568, 2019. ISSN 0002-8487 1548-8659. doi: 10.1002/tafs.10160.

43. V. Sbragaglia, C. Gliese, D. Bierbach, A. E. Honsey, S. Uusi-Heikkila, and R. Arlinghaus. Size-selective harvesting fosters adaptations in mating behaviour and reproductive allocation, affecting sexual selection in fish. Journal of Animal Ecology, 88(9):1343–1354, 2019. ISSN 0021-8790. doi: 10.1111/1365-2656.13032.

44. Tamal Roy, Kim Fromm, Valerio Sbragaglia, David Bierbach, and Robert Arlinghaus. Size selective harvesting does not result in reproductive isolation among experimental lines of zebrafish, danio rerio: Implications for managing harvest-induced evolution. Biology, 10(2):113, 2021. ISSN 2079-7737.

45. S. Uusi-Heikkilä, C. Wolter, T. Meinelt, and R. Arlinghaus. Size-dependent reproductive success of wild zebrafish *Danio rerio* in the laboratory. Journal of Fish Biology, 77(3):552–69, 2010. ISSN 0022-1112. doi: 10.1111/j.1095-8649.2010.02698.x.

46. K. O. Darrow and W. A. Harris. Characterization and development of courtship in zebrafish, *Danio rerio*. Zebrafish, 1(1):40–5, 2004. ISSN 1545-8547. doi: 10.1089/154585404774101662.

47. Alfonso Pérez-Escudero, Julián Vicente-Page, Robert C. Hinz, Sara Arganda, and Gonzalo G. de Polavieja. idtracker: tracking individuals in a group by automatic identification of unmarked animals. Nature Methods, 11(7):743–748, 2014. doi: 10.1038/nmeth.2994 https://www.nature.com/articles/nmeth.2994#supplementary-information.

48. S. Nakagawa and H. Schielzeth. Repeatability for gaussian and non-gaussian data: a practical guide for biologists. Biol Rev Camb Philos Soc, 85(4):935–56, 2010. ISSN 0006-3231. doi: 10.1111/j.1469-185X.2010.00141.x.

49. G. Polverino, D. Bierbach, S. S. Killen, S. Uusi-Heikkila, and R. Arlinghaus. Body length rather than routine metabolic rate and body condition correlates with activity and risk-taking in juvenile zebrafish *Danio rerio*. Journal of Fish Biology, 89(5):2251–2267, 2016. ISSN 0022-1112. doi: 10.1111/jfb.13100.

50. Pawel Romanczuk, Markus Bär, Werner Ebeling, Benjamin Lindner, and Lutz Schimansky-Geier. Active brownian particles. The European Physical Journal Special Topics, 202(1):1–162, 2012.

51. Ariana Strandburg-Peshkin, Colin R. Twomey, Nikolai W F Bode, Albert B. Kao, Yael Katz, Christos C. Ioannou, Sara B. Rosenthal, Colin J. Torney, Hai Shan Wu, Simon A. Levin, and Iain D. Couzin. Visual sensory networks and effective information transfer in animal groups. Current Biology, 23(17):R709–R711, 2013. ISSN 09609822. doi: 10.1016/j.cub.2013.07.059.

52. Nikolaus Hansen and Andreas Ostermeier. Adapting arbitrary normal mutation distributions in evolution strategies: The covariance matrix adaptation. Proceedings of IEEE International Conference on Evolutionary Computation, pages 312–317, 1996. doi: 10.1109/ICEC.1996.542381.

53. Nikolaus Hansen, Youhei Akimoto, and Petr Baudis. Cma-es/pycma on github. Zenodo, DOI:10.5281/zenodo.2559634, 2019.

54. Paolo Domenici. The scaling of locomotor performance in predator–prey encounters: from fish to killer whales. Comparative Biochemistry and Physiology Part A: Molecular & Integrative Physiology, 131(1):169–182, 2001. ISSN 1095-6433. doi: https://doi.org/10.1016/S1095-6433(01)00465-2.

55. Howard C Howland. Optimal strategies for predator avoidance: the relative importance of speed and manoeuvrability. Journal of Theoretical Biology, 47(2):333–350, 1974. ISSN 0022-5193.

56. Laurie Landeau and John Terborgh. Oddity and the ‘confusion effect’in predation. Animal Behaviour, 34(5):1372–1380, 1986. ISSN 0003-3472.

57. Parisa Rahmani, Fernando Peruani, and Pawel Romanczuk. Flocking in complex environments—attention trade-offs in collective information processing. PLOS Computational Biology, 16(4):e1007697, 2020. doi: 10.1371/journal.pcbi.1007697.

58. Nikolai W.F. Bode, Jolyon J. Faria, Daniel W. Franks, Jens Krause, and A. Jamie Wood. How perceived threat increases synchronization in collectively moving animal groups. Proceedings of the Royal Society B: Biological Sciences, 277(1697):3065–3070, 2010. ISSN 14712970. doi: 10.1098/rspb.2010.0855.

59. Pawel Romanczuk, Iain D. Couzin, and Lutz Schimansky-Geier. Collective motion due to individual escape and pursuit response. Physical Review Letters, 102(1):010602, jan 2009. ISSN 0031-9007. doi: 10.1103/PhysRevLett.102.010602.

60. Tamás Vicsek, András Czirók, Eshel Ben-Jacob, Inon Cohen, and Ofer Shochet. Novel type of phase transition in a system of self-driven particles. Physical Review Letters, 75(6):1226–1229, aug 1995. ISSN 0031-9007. doi: 10.1103/PhysRevLett.75.1226.

61. Jure Demšar, Charlotte K. Hemelrijk, Hanno Hildenbrandt, and Iztok Lebar Bajec. Simulating predator attacks on schools: Evolving composite tactics. Ecological Modelling, 304: 22-33, may 2015. ISSN 03043800. doi: 10.1016/j.ecolmodel.2015.02.018.

62. Randal S. Olson, David B. Knoester, and Christoph Adami. Evolution of swarming behavior is shaped by how predators attack. Artificial Life, 22(3):299–318, 2016. doi: 10.1162/ARTL_a_00206%M27139941.

63. Iain D. Couzin, Jens Krause, Nigel R Franks, and Simon A. Levin. Effective leadership and decision making in animal groups on the move. Nature, 13(4):513–516, 2005. ISSN 14607425. doi: 10.1038/nature03236.

64. Jolle W Jolles, Neeltje J Boogert, Vivek H Sridhar, Iain D Couzin, and Andrea Manica. Consistent individual differences drive collective behavior and group functioning of schooling fish. Current Biology, 27(18):2862–2868, 2017. ISSN 0960-9822.

65. Matthew M. G. Sosna, Colin R. Twomey, Joseph Bak-Coleman, Winnie Poel, Bryan C. Daniels, Pawel Romanczuk, and Iain D. Couzin. Individual and collective encoding of risk in animal groups. Proceedings of the National Academy of Sciences, 116(41):20556–20561, 2019. doi: 10.1073/pnas.1905585116.

66. Katja Enberg, Christian Jørgensen, Erin S. Dunlop, Øystein Varpe, David S. Boukal, Loïc Baulier, Sigrunn Eliassen, and Mikko Heino. Fishing-induced evolution of growth: concepts, mechanisms and the empirical evidence. Marine Ecology, 33(1):1–25, 2012. ISSN 1439-0485. doi: 10.1111/j.1439-0485.2011.00460.x.

67. Stephanie M. Carlson, Curry J. Cunningham, and Peter A. H. Westley. Evolutionary rescue in a changing world. Trends in Ecology & Evolution, 29(9):521–530, 2014. ISSN 0169-5347. doi: https://doi.org/10.1016/j.tree.2014.06.005.

68. Clémentine Renneville, Alexis Millot, Simon Agostini, David Carmignac, Gersende Maugars, Sylvie Dufour, Arnaud Le Rouzic, and Eric Edeline. Anthropogenic selection along directions of most evolutionary resistance. bioRxiv, page 498683, 2018. doi: 10.1101/498683.

69. Serinde J. van Wijk, Martin I. Taylor, Simon Creer, Christine Dreyer, Fernanda M. Rodrigues, Indar W. Ramnarine, Cock van Oosterhout, and Gary R. Carvalho. Experimental harvesting of fish populations drives genetically based shifts in body size and maturation. Frontiers in Ecology and the Environment, 11(4):181–187, 2013. ISSN 1540-9309. doi: 10.1890/120229.

70. Katja Enberg, Christian Jørgensen, Erin S. Dunlop, Mikko Heino, and Ulf Dieckmann. Implications of fisheries-induced evolution for stock rebuilding and recovery. Evolutionary Applications, 2(3):394–414, 2009. ISSN 1752-4571. doi: 10.1111/j.1752-4571.2009.00077.x.

71. Henry F. Wootton, Asta Audzijonyte, and John Morrongiello. Multigenerational exposure to warming and fishing causes recruitment collapse, but size diversity and periodic cooling can aid recovery. Proceedings of the National Academy of Sciences, 118(18):e2100300118, 2021. doi: 10.1073/pnas.2100300118.

72. Marion Claireaux, Christian Jørgensen, and Katja Enberg. Evolutionary effects of fishing gear on foraging behavior and life-history traits. Ecology and Evolution, 8(22):10711–10721, 2018. ISSN 2045-7758 (Print) 2045-7758 (Linking). doi: 10.1002/ece3.4482.

73. Christian Jørgensen and Rebecca E. Holt. Natural mortality: Its ecology, how it shapes fish life histories, and why it may be increased by fishing. Journal of Sea Research, 75:8–18, 2013. ISSN 1385-1101. doi: http://dx.doi.org/10.1016/j.seares.2012.04.003.

74. Christian Jørgensen and Oyvind Fiksen. Modelling fishing-induced adaptations and consequences for natural mortality. Canadian Journal of Fisheries and Aquatic Sciences, 67(7):1086–1097, 2010. ISSN 0706-652X. doi: 10.1139/F10-049.

75. Diana M. T. Sharpe and Andrew P. Hendry. Life history change in commercially exploited fish stocks: an analysis of trends across studies. Evolutionary Applications, 2(3):260–275, 2009. ISSN 1752-4571. doi: 10.1111/j.1752-4571.2009.00080.x.

76. Jeffrey A Hutchings. Influence of growth and survival costs of reproduction on atlantic cod, *Gadus morhua*, population growth rate. Canadian Journal of Fisheries and Aquatic Sciences, 56(9):1612–1623, 1999. ISSN 0706-652X.

77. Robert NM Ahrens, Carl J Walters, and Villy Christensen. Foraging arena theory. Fish and fisheries, 13(1):41–59, 2012. ISSN 1467-2979.

78. FAO. The state of world fisheries and aquaculture (sofia). sustainability in action. food and agriculture organization, rome, italy, pp. 244. Report, 2020.

79. Jeffrey A. Hutchings and Anna Kuparinen. Implications of fisheries-induced evolution for population recovery: Refocusing the science and refining its communication. Fish and Fisheries, 21(2):453–464, 2019. ISSN 1467-2960 1467-2979. doi: 10.1111/faf.12424.

80. Malin L. Pinsky, Anne Maria Eikeset, Cecilia Helmerson, Ian R. Bradbury, Paul Bentzen, Corey Morris, Agata T. Gondek-Wyrozemska, Helle Tessand Baalsrud, Marine Servane Ono Brieuc, Olav Sigurd Kjesbu, Jane A. Godiksen, Julia M. I. Barth, Michael Matschiner, Nils Chr. Stenseth, Kjetill S. Jakobsen, Sissel Jentoft, and Bastiaan Star. Genomic stability through time despite decades of exploitation in cod on both sides of the atlantic. Proceedings of the National Academy of Sciences, 118(15):e2025453118, 2021. doi: 10.1073/pnas.2025453118.

81. Jeffrey A. Hutchings and Anna Kuparinen. Throwing down a genomic gauntlet on fisheries-induced evolution. Proceedings of the National Academy of Sciences, 118(20):e2105319118, 2021. doi: 10.1073/pnas.2105319118.

82. Yang Xiang, Sylvain Gubian, Brian Suomela, and Julia Hoeng. Generalized simulated annealing for global optimization: The GenSA package. R J., 5(1):13–28, 2013. ISSN 20734859. doi: 10.32614/rj-2013-002.

83. Rainer Storn and Kenneth Price. Differential Evolution - A Simple and Efficient Heuristic for Global Optimization over Continuous Spaces. J. Glob. Optim., 11:341–359, 1997. doi: 10.1023/A:1008202821328.

